# Hydration conditions as a critical factor in antibiotic-mediated bacterial competition outcomes

**DOI:** 10.1101/2024.06.13.598809

**Authors:** Yana Beizman-Magen, Tomer Orevi, Nadav Kashtan

## Abstract

Antibiotic secretion plays a pivotal role in bacterial interference competition, yet the impact of environmental hydration conditions on such competition is not well-understood. Here, we investigate how hydration conditions affect interference competition among bacteria, studying the interactions between the antibiotic-producing *Bacillus velezensis* FZB42 and two bacterial strains susceptible to its antibiotics: *Xanthomonas euvesicatoria* 85-10 and *Pseudomonas syringae* DC3000. Our results show that wet-dry cycles significantly modify the response of the susceptible bacteria to both the supernatant and cells of the antibiotic-producing bacteria, compared to constantly wet conditions. Notably, *X. euvesicatoria* shows increased protection against both the cells and supernatants of *B. velezensis* under wet-dry cycles, while *P. syringae* cells become more susceptible under wet-dry cycles. In addition, we observed a reciprocal interaction between *P. syringae* and *B. velezensis*, where *P. syringae* inhibits *B. velezensis* under wet conditions. Our findings highlight the important role of hydration conditions in shaping bacterial interference competition, providing valuable insights into microbial ecology of water-unsaturated surfaces, with implications for applications such as biological control of plant pathogens and mitigating antibiotic resistance.

## Introduction

Bacteria, ubiquitous members of every ecosystem on our planet, play important roles in the maintenance and functioning of ecological processes^1–3^. These microorganisms do not live in isolation, but within complex communities, where interactions among different species are common^4,5^. The intricate interactions among bacterial species play a large role in shaping community composition and dynamics ^4,6–10^. Such interactions within microbial communities underlie important features of microbial communities such as diversity and stability^9,10^. Deciphering the nature and outcomes of these interactions is crucial for understanding the complexities of microbial ecology and its impact on ecological systems and processes^4,9^.

Bacterial interspecies competition is a common interaction that can take an active or passive forms^11–13^. The passive form involves competition for resources such as nutrients or space, whereas active competition involves one species secreting compounds that actively inhibit the growth or kill other species. This type of competition is called interference competition^11,13,14^. While interference competition commonly occurs naturally, it can also be artificially induced, as in the application of bacterial biological control against plant pathogens^12,15^, or in bioremediation practices^16^.

The outcome and dynamics of most interspecific interactions among bacteria can be influenced by chemical, physical and biological characteristics of the environment. Factors such as nutrient availability, temperature, pH, and surface type, significantly affect these interactions^17–22^. Among these factors, hydration conditions – which refer to the availability of water or moisture, ranging from extremely dry to fully saturated environments – play a pivotal role in many terrestrial microbial habitats. Hydration conditions are governed by the complex interplay of the presence of water or vapor, relative humidity, temperature, and the chemical and physical characteristics of surfaces^23–29^. Typically, in environments that are not constantly saturated with water, hydration conditions are not static, but rather fluctuate dynamically, often characterized by recurring wet-dry cycles^30–32^.

During the ‘dry’ phases of wet-dry cycles, surfaces can retain Microscopic Surface Wetness (MSW ^25,30,33^). MSW can take the form of microscopic droplets or thin liquid films, often invisible to the naked eye. A primary factor for MSW formation and retention on surfaces is the presence of deliquescent substances, such as highly hygroscopic salts. For example, on leaf surfaces a major source of salts comes from atmospheric aerosols present ubiquitously across plant foliage globally^25,30^. These salts can form, or retain, microscopic wetness when the relative humidity is above the deliquescence, or efflorescence, points^25,30^. MSW is prevalent on both biotic and abiotic surfaces^33–36^ and has been shown to often form around bacterial aggregates and other microorganisms^25,33^. Moreover, MSW has been found to protect bacteria and other microorganisms from desiccation and affect their survival rates^25,33,37^. Microscopic wetness has been shown to significantly impact bacterial life and ecology, affecting mobility, communication, horizontal gene transfer^38^, and spatial organization during colonization of surfaces^25,39^.

In a recent study we have revealed that wet-dry cycles with MSW can protect bacteria from major antibiotic classes^40^. We found that under wet-dry cycles, bacterial response to major antibiotics classes, markedly differs than their reaction under constantly wet conditions. Under wet-dry cycles with a period of MSW conditions (at the ‘dry’ period), bacteria showed increased protection from diverse antibiotic classes, mostly antibiotic classes that are effective against actively growing cells. Through a combination of experiments and computational modeling, we suggested four mechanisms, operating at different phases of a wet-dry cycles, that lead to increased protection from antibiotics under wet-dry cycles: (1) cross-protection due to high salt concentrations, (2) ‘tolerance by slow growth’, (3) deactivation of antibiotics by the physicochemical conditions associated with drying and MSW, and (4) ‘tolerance by lag’ during rewetting^40^. Given that hydration conditions and the presence of MSW notably affect bacterial responses to antibiotics, we hypothesize that these factors may also exert an impact on antibiotic-mediated bacterial interference competition outcomes.

To explore how wet-dry cycles with MSW affect interference competition among bacteria, we designed an experimental setup that enables the incubation of bacterial cultures within droplets of various sizes, maintained either wet (at 100% relative humidity) or subjected to wet-dry cycles (at 85% relative humidity) featuring MSW periods. We hypothesize that the response of antibiotic-sensitive bacteria, to antibiotic-producing bacteria, under a wet-dry cycle with MSW, can be significantly different from their response under constantly wet conditions. For our investigation, we selected the biological control agent *Bacillus velezensis* FZB42 (*Bv*FZB42)^41^, renowned for its ability to produce a broad spectrum of antimicrobial compounds, including several antibiotics^41^. We also chose two well-studied plant foliar pathogens: *Xanthomonas euvesicatoria pv. Vesicatoria 85-10 (Xee*85-10)^42,43^, and *Pseudomonas* syringae DC3000 (*Pst*DC3000) ^44,45^. We incubated cells of *Xee*85-10 or *Pst*DC3000 in droplets of different sizes, either alone or in combination with *Bv*FZB42 supernatants, cells, or both, under wet-dry cycle conditions and compared the outcomes (based on quantifying colony forming units) to those under constantly wet conditions. Cell numbers and competition outcomes were then determined by.

## Results

### Inhibition of *Xee*85-10 and *Pst*DC3000 by *Bv*FZB42 supernatants and cells under standard lab assays

Initially, we sought to determine the inhibitory capabilities of *Bv*FZB42 cells and/or their supernatants (see Methods) against *Xee*85-10 and *Pst*DC3000. Through two established assays – an inhibition zone assay and a MIC assay (specifically for supernatants) – we observed that both the cells and the supernatants of *Bv*FZB42 exerted a significant antagonistic effect against both *Xee*85-10 and *Pst*DC3000. This effect was evident in the results of inhibition zone assays (Fig. 1A), as well as Minimal Inhibitory Concentration (MIC) assays (as depicted in Fig. 1B).

**Figure 1.**
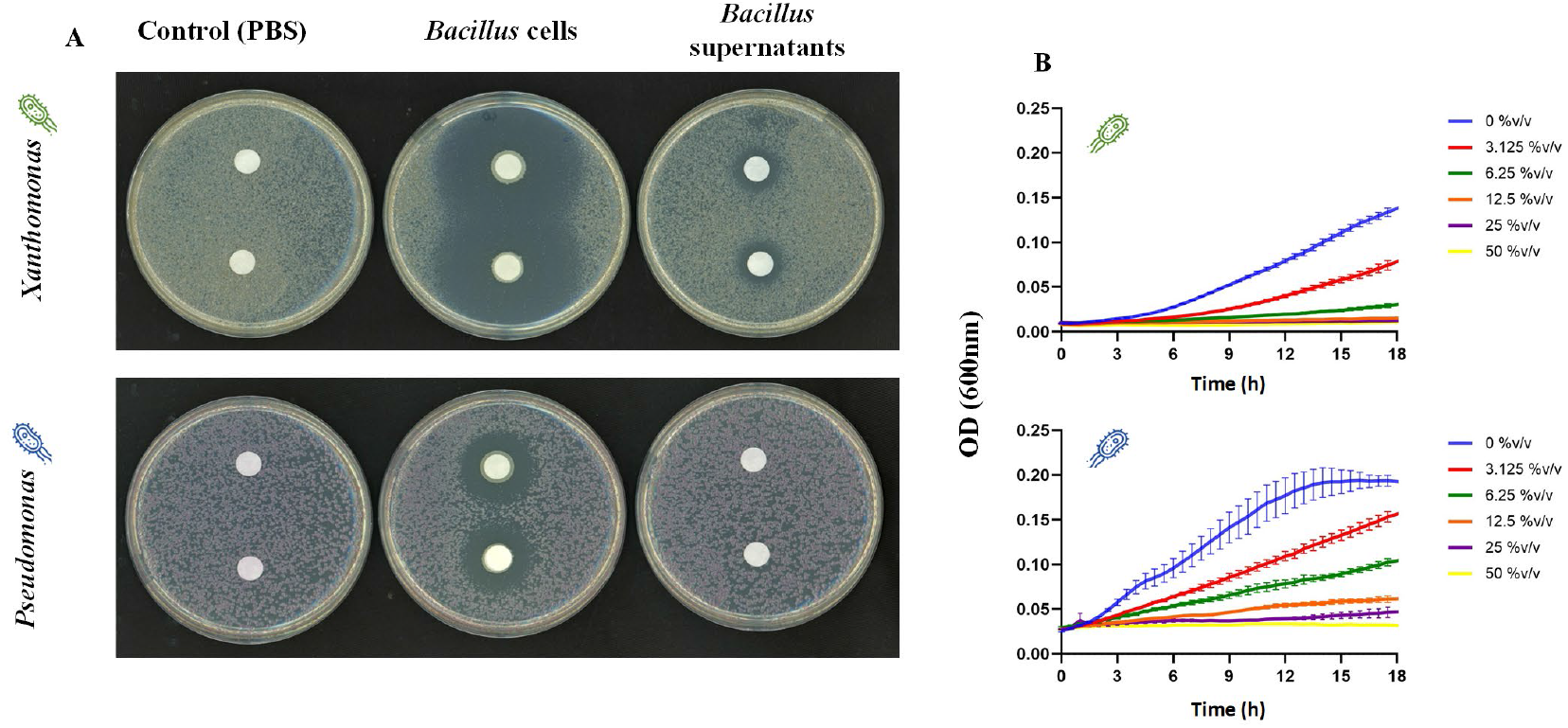
Effectiveness of *Bv*FZB42 cells or supernatants against *Xee*85-10 and *Pst*DC3000. (A) Inhibition zone assay of *Xee*85-10 (top) and *Pst*DC3000 (bottom) by *Bv*FZB42 supernatants (100%, right), cells (OD600=0.1, center) +cells (center), live bacterial cells (right) and control (PBS, left). (B) determination of minimal inhibitory concentration (MIC) of *Xee*85-10 (top) and *Pst*DC3000 (bottom), respectively, by *Bv*FZB42 supernatants.

Notably, *Pst*DC3000 was inhibited in 50% v/v concentration of *Bv*FZB42 supernatants (meaning when the volume consisted of 50% *Bv*FZB42 supernatants), while *Xee*85-10 required a lower concentration of 12.5% v/v for inhibition.

### Design of an experimental setup for bacterial culturing under constant wetness or wet-dry cycle conditions

To explore the effects of hydration conditions on bacterial interference competition, our experimental setup involved culturing bacteria within drying droplets, as detailed in Methods. A key variable influencing the drying dynamics of these droplets is their initial size. To comprehensively investigate the range of drying dynamics and corresponding ‘drying times’, we utilized droplets varying in volume from 1 µL to 100 µL. This range was selected to represent a spectrum of drying durations under controlled conditions (under moderate relative humidity). Our observations confirmed that droplet volume dictates the time required to reach microscopic surface wetness (MSW) conditions, with short drying duration of ∼1 h in the smallest droplets and longer drying times of ∼8 h in the largest droplets (see Supp. Fig. S1).

To assess and compare the influence of continuous wet conditions versus a 24-hour wet-dry cycle, we maintained a subset of droplets at a constant humidity of 100% RH, ensuring they remained static (i.e. kept hydrated) throughout the experiment, while others were subjected to a cycle of drying (under moderate relative humidity, RH=85%) until stable MSW formed, followed by rewetting. This design allowed us to closely monitor and analyze the effects of hydration dynamics on the competition outcome (by CFU plating at the beginning and end of the experiments) under these distinct environmental conditions.

### *Xee*85-10 is less affected by *Bv*FZB42 supernatants under wet-dry cycle compared to constantly wet conditions

First, we established a baseline by evaluating *Xee*85-10’s growth over 24 hours without the presence of *Bv*FZB42’s supernatants. Under constantly wet conditions, *Xee*85-10 exhibited a growth increase, of up to 2 log10 CFU in the larger droplet volumes, indicating robust growth capacity in an environment free from desiccation stress (one-way ANOVA, p<0.05, Fig. 2A; Supp. Fig. S2). In contrast, under a wet-dry cycle, a different pattern emerged. In all droplet volumes except for the largest (100µl), we observed a reduction in *Xee*85-10 CFUs, which suggests that the bacterium’s survival was adversely affected by the cyclic drying and rewetting process. This impact was more pronounced in smaller droplets, the faster drying dynamics of the smaller droplets reduced cell viability (one-way ANOVA, p<0.05, Fig. 2; Supp. Fig. S2, S3).

**Figure 2.**
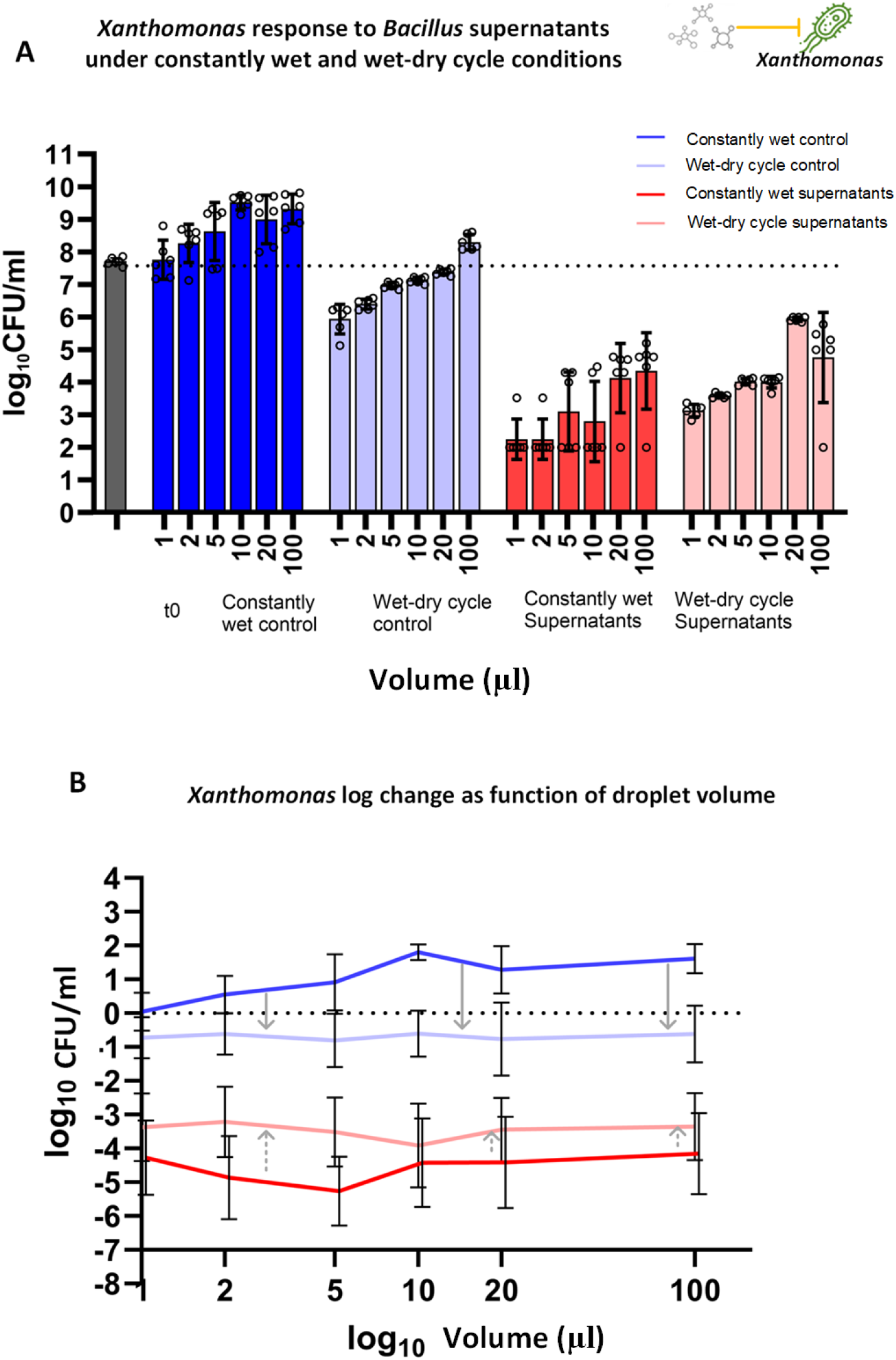
*Xee*85-10 response to *Bv*FZB42 supernatants under wet-dry cycle and constantly wet conditions. **(A)** Log 10 CFU of *Xee*85-10 under constantly wet and wet-dry cycle conditions with or without *Bv*FZB42 supernatants, at t=24 h. The most left grey bar represents CFU/ml at t=0 h. Bars and error bars represent mean ± SE CFU/ml. Black circles represent technical replicates. CFU measurements were conducted as described in Methods. One-way ANOVA was performed to compare the means of all conditions with each other (Supp. Fig. S2). **(B)** Log change in CFU/ml (log10 CFU at t=24 h minus log10 CFU at t=0 h), as a function of droplet size (X-axis in log scale), in the presence and absence of supernatants. Lines and error bars represent mean ± SD. Linear regression and Spearman correlation were performed for statistical assessment (Supp. Fig. S3). Grey arrows represent the changes between wet-dry cycle and constantly wet conditions in control (solid arrows) and with the addition of supernatants (dashed arrows).

Upon addition of *Bv*FZB42 supernatants at a concentration of 30% v/v (Methods), a notable reduction in CFU, of ∼3-5 logs, was observed under constantly wet conditions, aligning with expectations set by previous inhibition assays (Fig. 1) and given that the supernatant concentration exceeded the MIC of 12.5% v/v (one-way ANOVA, p<0.0005, Fig. 2A; Supp. Fig. S2). Under wet-dry cycle, all drop volumes demonstrated a significant CFU reduction from the baseline (t = 0 h) (ANOVA, p<0.05, Fig. 2A; Supp. Fig. S2).

Notably, the CFU reduction was less pronounced under wet-dry cycles when supernatants were introduced, compared to the scenario without supernatant (ANOVA, p<0.05, Fig. 2A, B; Supp. Fig. S2). These results show that *Xee*85-10’s is more protected from *Bv*FZB42 supernatants under wet-dry conditions relative to constant wetness.

### The effectiveness of *Bv*FZB42 supernatants against *Pst*DC3000 was markedly more pronounced under wet-dry cycle as opposed to constantly wet conditions

Without supernatants, *Pst*DC3000, CFU counts increased under constantly wet conditions and decreased under wet-dry cycles, with larger decrease in smaller droplets (Fig. 3; Supp. Figs. S4, S5), similarly to the response pattern of *Xee*85-10. This pattern indicates a general decline in bacterial viability under the stress of wet-dry cycles, though with improved survival rate in larger droplets (one-way ANOVA, p<0.05, Fig. 3; Supp. Fig. S4).

**Figure 3.**
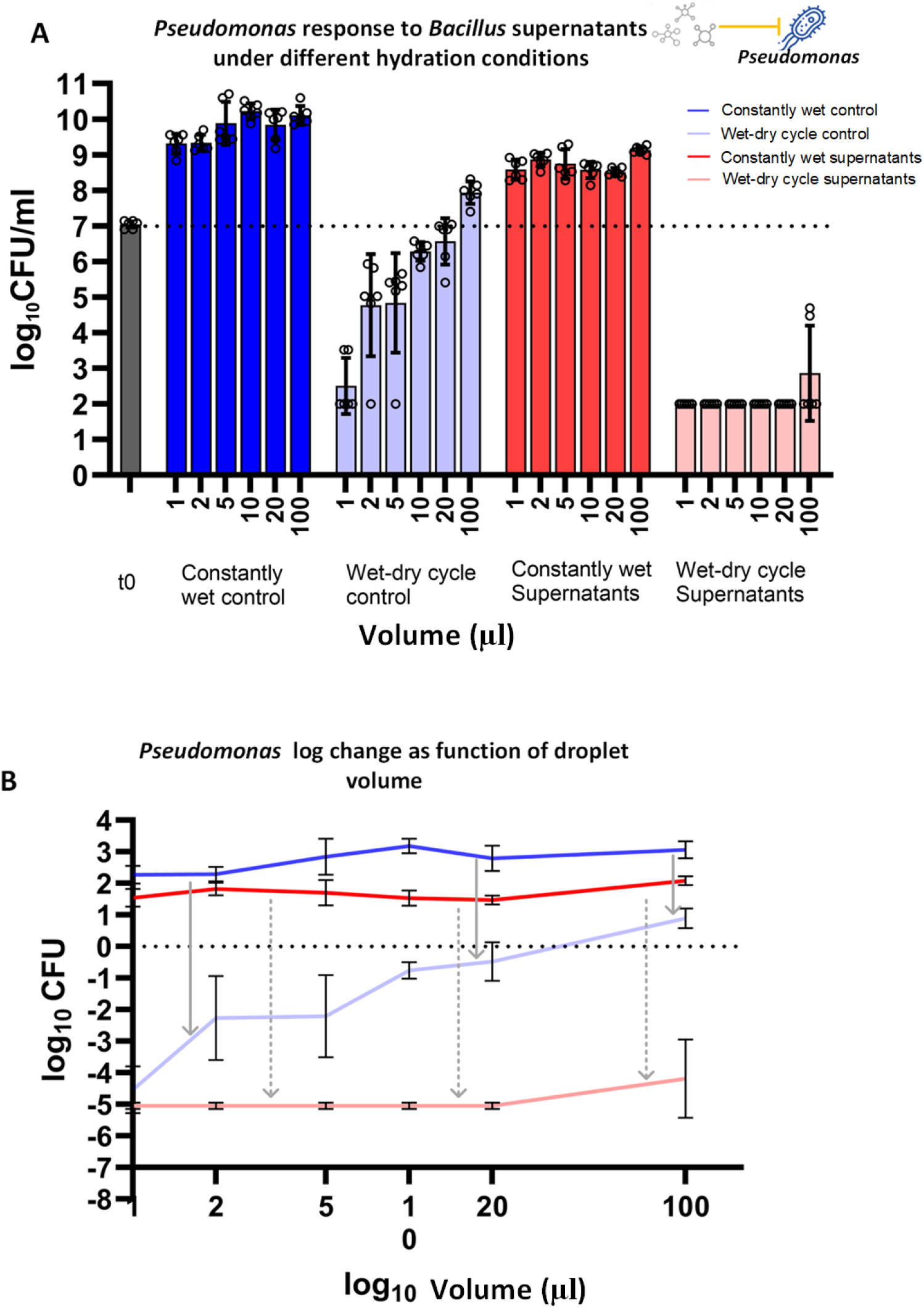
*Pst*DC3000 response to *Bv*FZB42 supernatants under wet-dry cycle and constantly wet conditions. **(A)** Log 10 CFU of *Pst*DC3000 exposed to *Bv*FZB42 supernatants under constantly wet and wet-dry cycle conditions, at t=24 h. The most left grey bar represents CFU/ml at t=0 h. Bars and error bars represent mean ± SE CFU/ml. Black circles represent technical replicates. CFU measurements were conducted as described in Methods. One-way ANOVA was performed to compare the means of all conditions with each other (Supp. Fig. S4). **(B)** Log change in CFU/ml (log10 CFU at t=24 h minus log10 CFU at t=0 h), as a function of droplet size (X-axis in log scale), in the presence and absence of supernatants. Lines and error bars represent mean ± SD. Linear regression and Spearman correlation were performed for statistical assessment (Supp. Fig. S5). Grey arrows represent the changes between wet-dry cycle and constantly wet conditions in control (solid arrows) and with the addition of supernatants (dashed arrows).

Introducing *Bv*FZB42 supernatants at a 30% v/v concentration (an equal concentration to the treatment for *Xee*85-10) elicited a distinct response in *Pst*DC3000. Under constant wetness, *Pst*DC3000’s CFU levels still increased from the baseline, as could be anticipated, given the supernatant concentration was below the MIC for *Pst*DC3000 (30% v/v versus an MIC of 50% v/v) (Fig. 4A; Supplementary Fig. S4). Intriguingly, the wet-dry cycle conditions triggered a dramatic CFU reduction across all droplet volumes, much more than under constant wet conditions (one-way ANOVA, p<0.0001, Fig. 3A; Supp.Fig. S4), as well as in comparison to wet-dry cycle without supernatants (one-way ANOVA, p<0.0001,Fig. 3A; Supp. Fig. S4). Thus, *Pst*DC3000 cells were much more affected by *Bv*FZB42 supernatants under a wet-dry cycle compared to constantly wet conditions.

**Figure 4.**
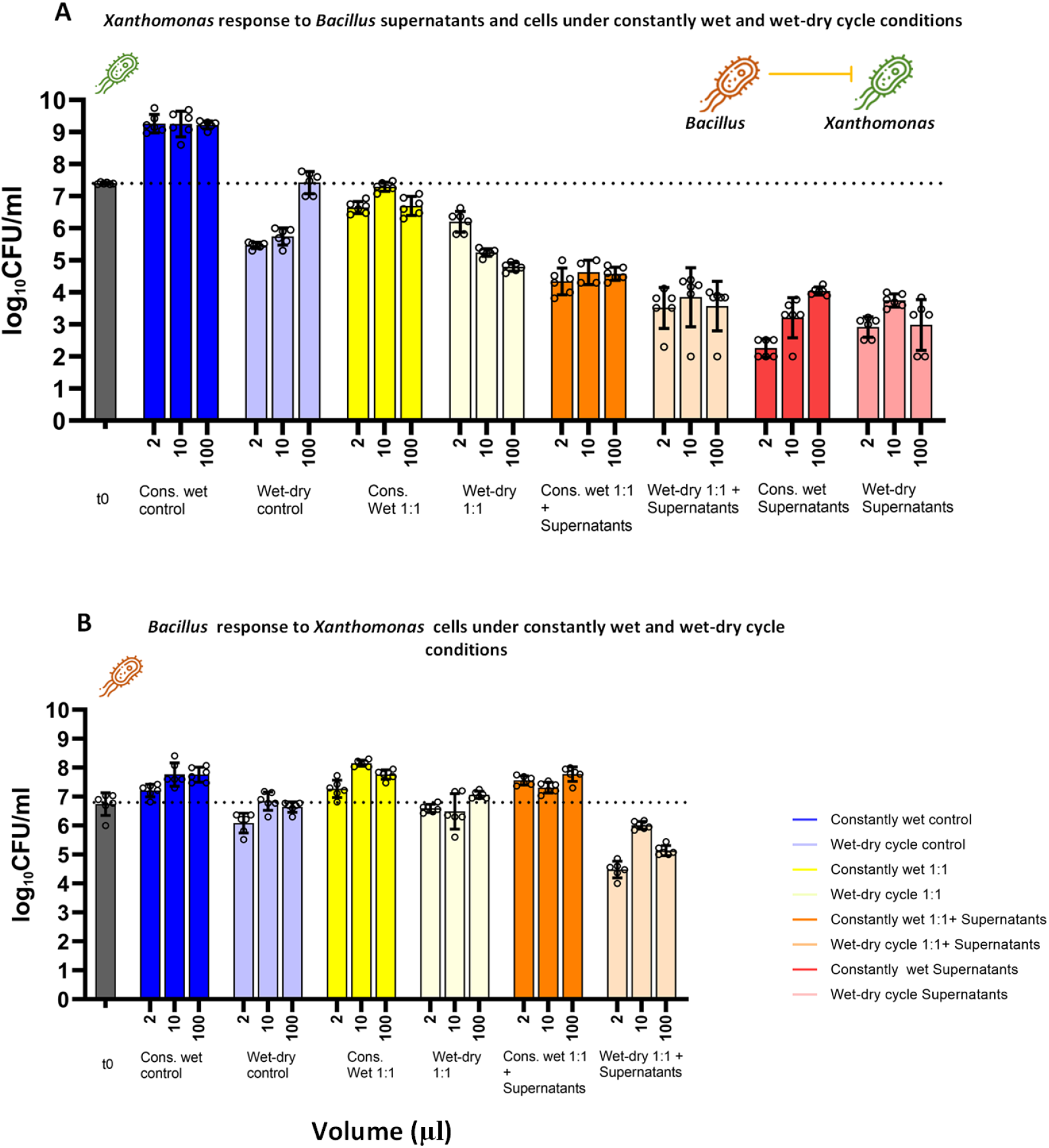
Co-culture experiments of *Xee*85-10 and *Bv*FZB42 under constantly wet and wet-dry cycle conditions. **(A)** CFU/ml of *Xee*85-10 at t=24 h under both constantly wet and wet-dry cycle conditions across different co-culture scenarios and controls and at 3 different droplet volumes. Left most grey bar represent CFU/ml at t=0 h. Bars and error bars represent mean ± SE CFU/ml. Black circles represent technical replicates. One-way ANOVA was performed to compare the means between conditions and droplet volumes with each other (Supp. Figs. S6, S8). (**B)** Same as in (A) but for *Bv*FZB42 (CFU/ml at t= 24 h).

### Co-culture experiments representing two ecological scenarios

Next, we introduced an additional layer of complexity by performing co-culture experiments of the sensitive bacteria together with cells of the antibiotic-producing bacteria, *Bv*FZB42, with or without its supernatants. We employed two types of co-culture experiments; each represent a different ecological scenario. In both scenarios there were similar concentration of both *Bv*FZB42 and the sensitive bacteria at t =0 h (∼10^7 CFU/ml). In the first, cells from *Bv*FZB42 and the pathogen (*Xee*85-10 or *Pst*DC3000) were co-cultured at an initial 1:1 cell ratio, simulating their concurrent arrival in a habitat (we term this scenario 1:1). The second scenario included both *Bv*FZB42 cells and its supernatants, resembling a situation where *Bv*FZB42 was already established and producing antibiotics before the pathogen’s arrival (we term this scenario 1:1 plus supernatants; 1:1 cell ratio of both species plus supernatants at t=0). This setup also aligns with biological control practices, such as foliar spraying, where both live microbial agents and their products are applied together^46–48^.

### *Xee*85-10 is less affected by *Bv*FZB42 cells under a wet-dry cycle, with and without supernatants

Under constantly wet conditions, there was a decrease in *Xee*85-10 CFUs when *Bv*FZB42 cells were added with, or without, their supernatants (Fig. 4A). Co-culturing *Xee*85-10 with *Bv*FZB42 cells without supernatants showed a modest decrease in *Xee*85-10 CFUs under wet conditions, and a more significant reduction under wet-dry conditions. Adding supernatants to the mix increased the CFU reduction further, under both conditions. Interestingly, when comparing the effects of constant wetness to wet-dry cycles with supernatants, CFU numbers were similar or slightly higher in the wet-dry cycle with supernatants only (Fig. 4A, Supp. Fig. S6). This observation reinforces the finding that *Xee*85-10 is less impacted by *Bv*FZB42 under wet-dry cycles.

The pronounced protection of *Xee*85-10 from cells and supernatants of *Bv*FZB42, under a wet-dry cycle, as opposed to constantly wet conditions, is in agreement with suspended liquid co-cultures in microwells (quantified via plate reader, see Methods). In such microwell co-cultures, *Xee*85-10 growth was completely inhibited by either *Bv*FZB42 cells, supernatants, or both (Supp. Fig. S7). However, in our droplet-based co-cultures, whether under constant wetness or wet-dry cycles, *Xee*85-10 remained viable after 24 h.

*Bv*FZB42, however, appeared to be unaffected by the presence of *Xee*85-10. Under both constantly wet and wet-dry cycle conditions, CFU counts were comparable in the mono-culture and co-culture setups (control, 1:1, and 1:1 plus supernatants). Under constantly wet conditions, there was a significant increase in *Bv*FZB42 CFU counts (one-way ANOVA, p<0.0001, Fig. 4B, Supp. Fig. S8). During a wet-dry cycle, a slight reduction in *Bv*FZB42 CFU counts was observed, which was more pronounced when supernatants were added (1:1 plus supernatants) (Fig. 4B). These results indicate that *Bv*FZB42’s growth, or survival, is not significantly affected by the presence of *Xee*85-10 in most conditions.

To enhance our understanding of the interactions between these two bacterial species, we have supplemented our previous data with area plots (Fig. 5, Supp. Fig. S9). These plots offer a visual representation of changes in the abundance of each species within a competitive context. Under constantly wet conditions, *Xee*85-10 was generally outcompeted by *Bv*FZB42, particularly in the presence of supernatants with a clear decrease in *Xee*85-10’s abundance. During wet-dry cycles, the growth of both bacteria was inhibited regardless of supernatant presence (Fig. 5). In most scenarios, *v*FZB42 was the dominant competitor after 24 hours. However, in the 1:1 scenario within small 2 µL droplets, *Xee*85-10 managed to outcompete *Bv*FZB42 (Supp. Fig. S9).

**Figure 5.**
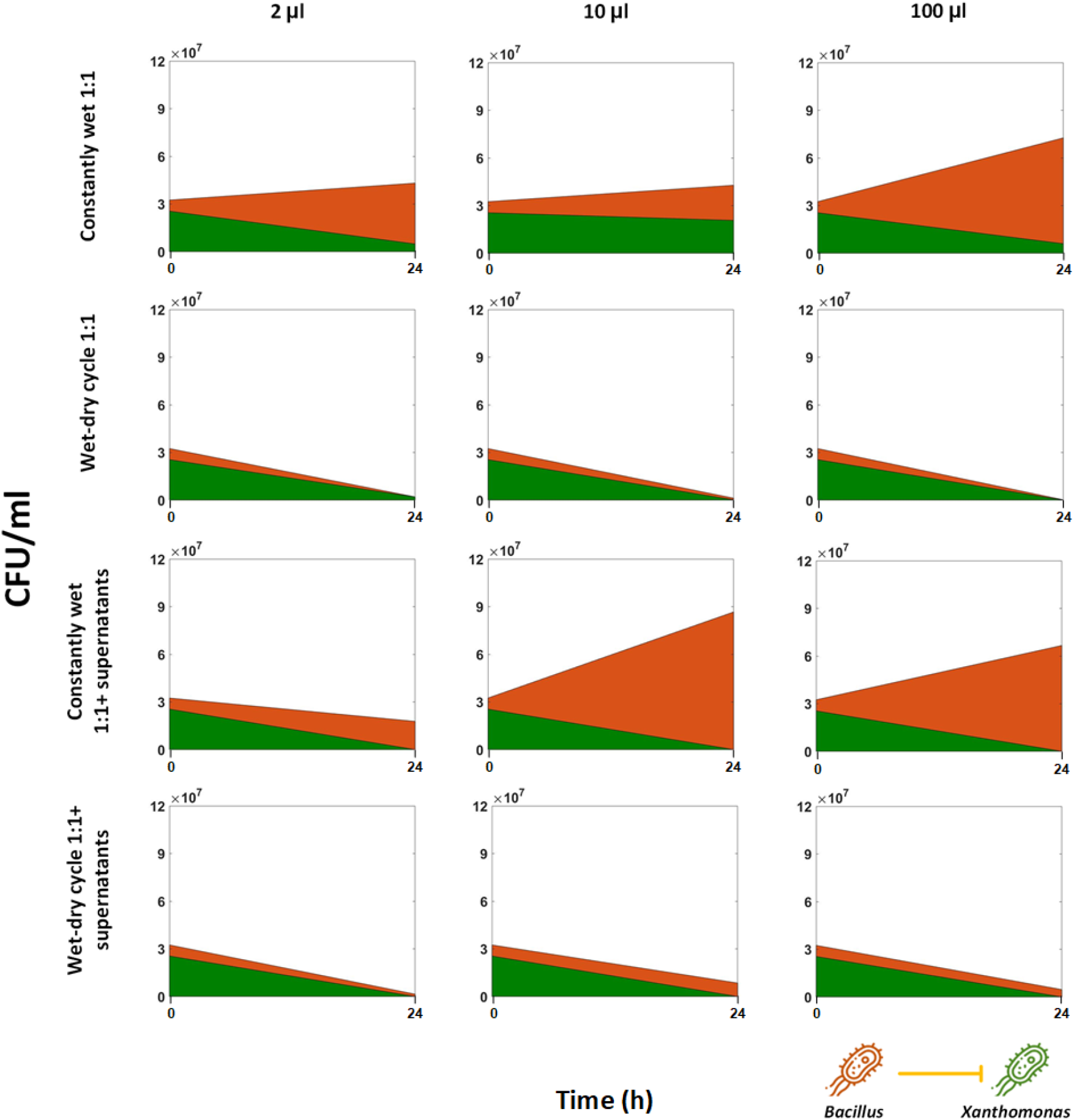
Competition dynamics of *Xee*85-10 and *Bv*FZB42 co-culture in droplets. The graph shows stacked area plots in droplet co-cultures (1:1 and 1:1+ supernatants). The green fraction represents *Xee*85-10, and the orange fraction represents *Bv*FZB42. Note the plot is based on only two time points (at t=0 h and t=24 h).

In summary, our findings suggest that *Bv*FZB42 inhibits *Xee*85-10 primarily due to the antibiotic compounds it secretes (present in its supernatants), yet this inhibitory effect is reduced under wet-dry cycles.

### The response of *PstDC3000* to *BvFZB42* cells and supernatants under wet-dry cycle differed from that of *Xee85-10*

Under constantly wet conditions, *Pst*DC3000 exhibited a variable response when co-cultured with *Bv*FZB42. A moderate reduction in *Pst*DC3000 CFUs was observed in 1:1 co-culture with *Bv*FZB42 cells alone, which became more pronounced when both *Bv*FZB42 cells and their supernatant were present (Fig. 6A, Supp. Fig. S10). Remarkably, consistent with experiments using only supernatants (Fig. 3), under wet-dry cycles, there was a complete inhibition of *Pst*DC3000(resulting in zero observed CFU) in co-culture with *Bv*FZB42, regardless of the presence of supernatants. The response of *Bv*FZB42 to interaction with *Pst*DC3000 in co-culture differed significantly from that with *Xee*85-10. Under both constantly wet and wet-dry conditions, *Bv*FZB42 experienced significant inhibition, with the exception being under constantly wet conditions with supernatants in 10 and 100 µL droplets (one-way ANOVA, p<0.005, Fig. 6B; Supp. Fig. S11).

**Figure 6.**
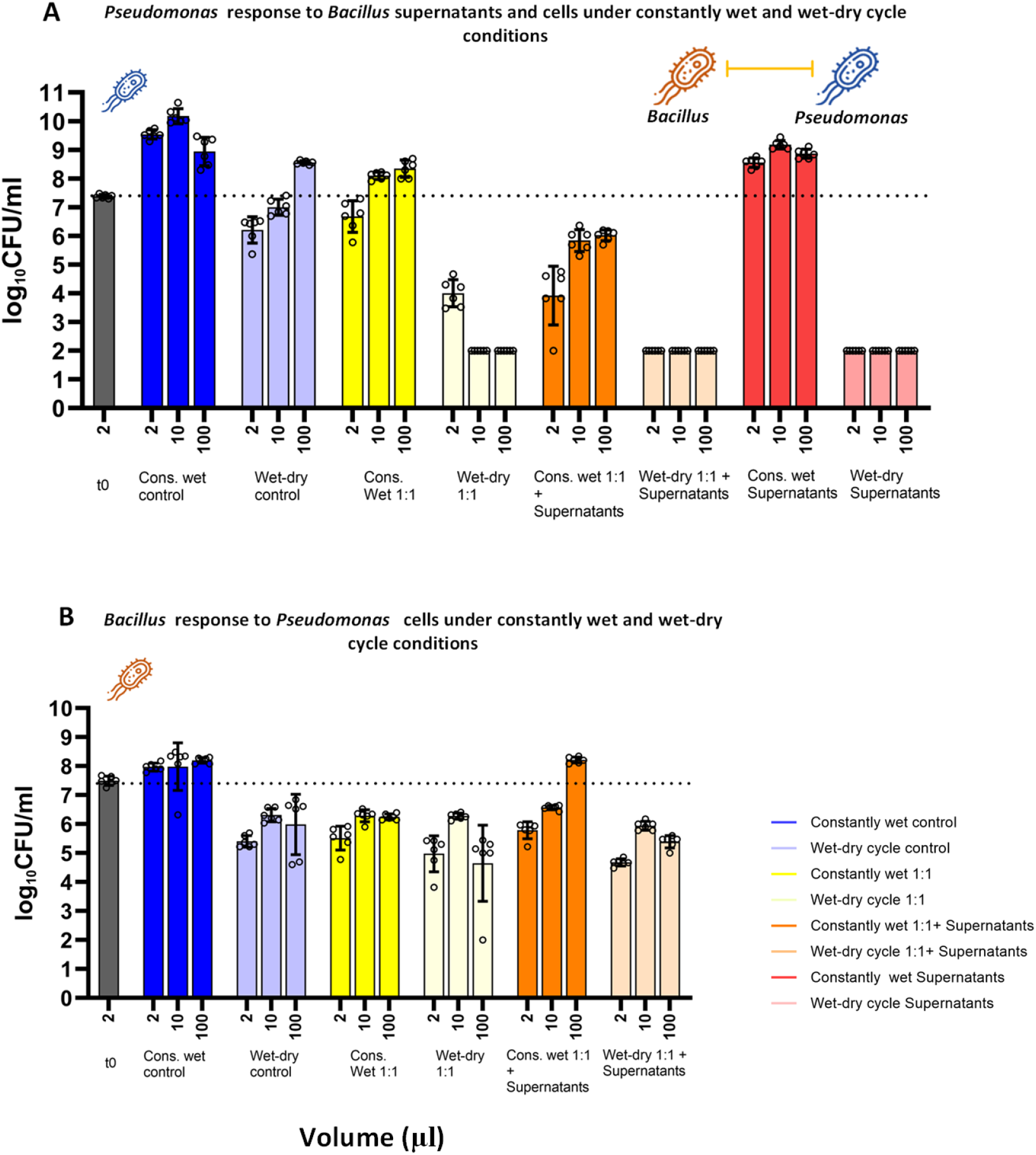
Co-culture experiments of *Pst*DC3000 and *Bv*FZB42 under constantly wet and wet-dry cycle conditions. **(A)** CFU/ml of *Xee*85-10 at t=24 h under both constantly wet and wet-dry cycle conditions across different co-culture scenarios and controls and at 3 different droplet volumes. Left most gray bar represent CFU/ml at t=0 h. Bars and error bars represent mean ± SE CFU/ml. Black circles represent technical replicates. One-way ANOVA was performed to compare the means between conditions and droplet volumes with each other (Supp. Figs. S10, S11). (**B)** Same as in (A) but for *Bv*FZB42 (CFU/ml at t=24 h).

These findings, revealing complete inhibition of *Pst*DC3000 under wet-dry cycles, were also very different from the results from co-cultures in suspended liquid cultures in micro wells, where *Pst*DC3000 remained viable in the presence of both *Bv*FZB42 cells and supernatants (1:1 ratio) (Supp. Fig. S12), mirroring the outcomes observed under constantly wet conditions in droplet experiments.

Under suspended liquid conditions, consistent with the droplet experiment, *Bv*FZB42 was inhibited. These observations further suggest that *Pst*DC3000 exerts an inhibitory effect on *Bv*FZB42, indicating a mutual inhibition between these two bacterial species (Supp. Figs. S12C, S13).

The dynamics of competition between *Bv*FZB42 and *Pst*DC3000 in co-culture experiments reveal varied outcomes, as illustrated by the area plots (Fig. 7). Consistent with the above results, under constantly wet conditions in co-culture without supernatant, *Pst*DC3000 took over the population. This outcome was reversed in the large drops (100 µL) when supernatants were added, where *Bv*FZB42 outcompeted *Pst*DC3000. In scenarios involving wet-dry cycles, regardless of supernatant presence, both *Pst*DC3000 and *Bv*FZB42 experienced a decline in CFUs. However, despite this reduction, the relative fraction of *Bv*FZB42 was larger (Supp. Fig. S14). These results highlight the significant impact of wet-dry cycles on the competition dynamics between these two bacterial species.

**Figure 7.**
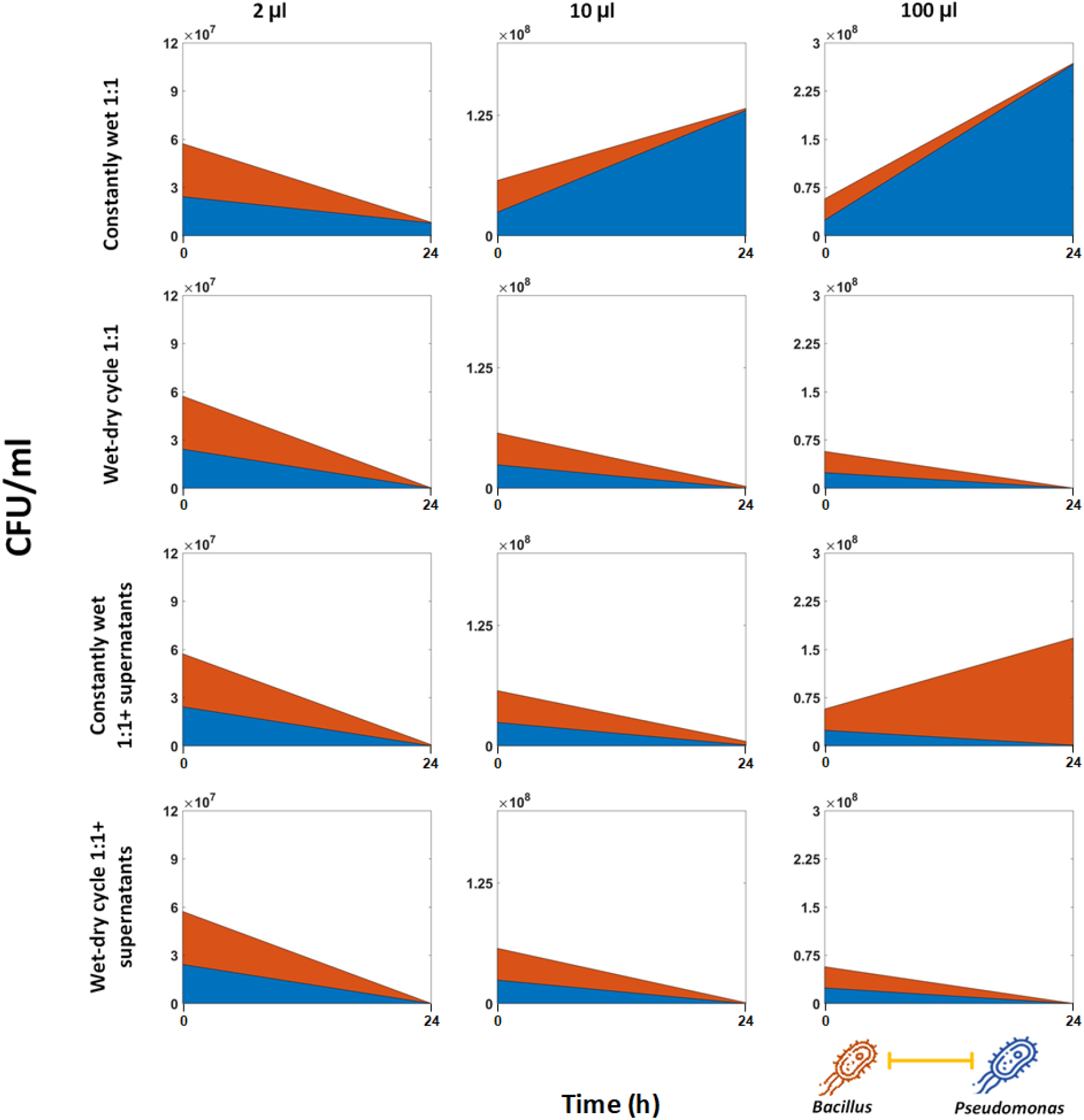
Competition dynamics of *PstDC3000*and *Bv*FZB42 co-culture in droplets. The graph shows stacked area plots in droplet co-cultures (1:1 and 1:1+ supernatants). The blue fraction represents *PstDC3000*, and the orange fraction represents *Bv*FZB42. Note the plot is based on two time points (at t=0 h and t=24 h).

### Discussion

In this study, we explored the interactions between the antibiotic-producing bacteria *Bv*FZB42 and two susceptible bacterial strains – *Xee*85-10 and *Pst*DC3000, under different hydration conditions, including constant wetness and wet-dry cycles. Our findings reveal that hydration conditions play a crucial role in determining the outcome of bacterial interference competition and the effectiveness of antibiotic-mediated inhibition. The differential responses of *Xee*85-10 and *Pst*DC3000 to *Bv*FZB42 cells and supernatants containing antibiotic compounds, highlight the potential for environmental factors, such as hydration conditions studied here, to modulate these interactions.

Our findings demonstrate how hydration condition critically influence the dynamics of antibiotic-mediated interference competition among bacteria, with a potential to either diminish or enhance the inhibitory effect (see visual summary in Fig. 8). Notably, under constant wet conditions, *Xee*85-10 experienced inhibition by *Bv*FZB42, yet it showed increased resilience against *Bv*FZB42’s supernatants during wet-dry cycles. *Pst*DC3000, in contrast, showed relative insensitivity to *Bv*FZB42’s supernatants under constantly wet environments, showed complete inhibition under wet-dry cycles. This variation in competitive outcomes across different hydration scenarios highlights the importance of considering environmental conditions when predicting competition results, rather than relying only on standard lab assays in suspended liquid or on agar plates.

**Figure 8.**
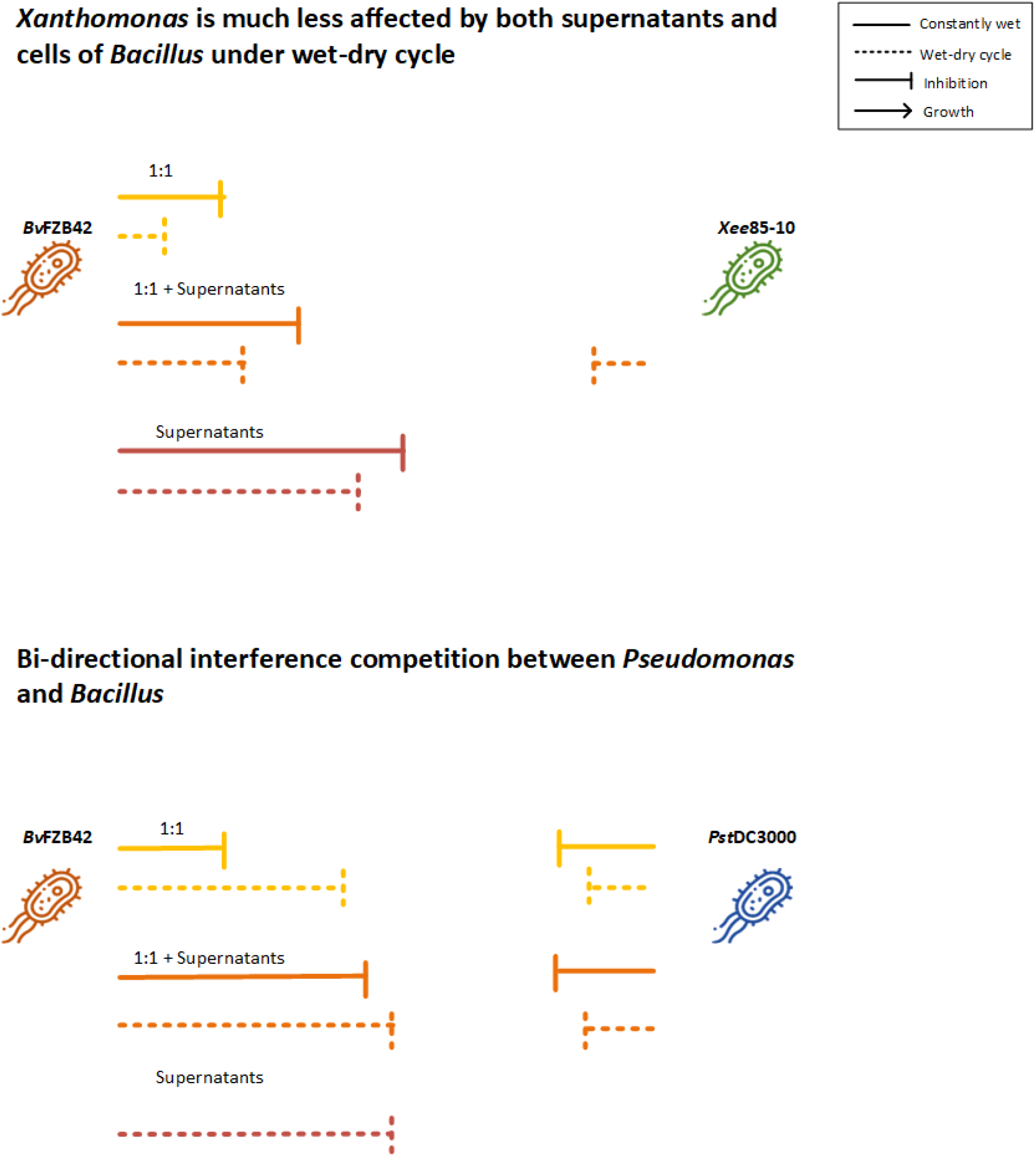
Graphical summary of the impact of hydration conditions on the interactions between *Bv*FZB42 and *Xee*85-10 or *Pst*DC3000. Top panel: *Xee*85-10 cells are less susceptible to *Bv*FZB42 cells and supernatants in wet-dry cycles compared to constantly wet conditions. The length of the inhibition arrow represents the inhibition strength. Bottom panel: *Pst*DC3000 is highly susceptible to *Bv*FZB42 supernatants in wet-dry cycle conditions but not under constantly wet conditions. The interaction is bi-directional as *Pst*DC3000 also inhibit *Bv*FZB42.

In a previous study, we showed that under wet-dry cycles bacteria can be more protected from diverse antibiotic classes^40^ .This phenomenon may explain the increased protection of *Xee*85-10 from the antibiotics produced by *Bv*FZB42 under wet-dry cycles. Other studies that explored the impact of *Bacillus velezensis* on *Xanthomonas* strains have demonstrated its effectiveness, however they were performed on other experimental setups, without an attempt to understand the effect of hydration conditions^49–52^. *Bv*FZB42 is known to produce a diverse array of antimicrobial compounds, including two antibiotics, difficidin and bacilysin, which have been proven effective against *Xanthomonas* strains ^52^. Difficidin, a macrolide, inhibits protein synthesis by binding to ribosomal subunits^53^, and may also compromise cell membranes^54^. Bacilysin, on the other hand, disrupts peptidoglycan biosynthesis through anticapsin, impacting bacterial growth in a similar way to beta-lactam antibiotics^55,56^. The mechanisms of these two antibiotics suggest that the increased protection of *Xee*85-10 could stem from a decreased susceptibility due to the slow or halted growth induced by drying, coupled with the low metabolic state triggered by the stresses of MSW conditions. This aligns with previous findings on the enhanced protection from beta-lactams under wet-dry cycles^40^. Additionally, the physicochemical conditions associated with MSW, such as elevated salt concentrations, increased levels of reactive oxygen species (ROS), and altered pH, may contribute to the degradation or inactivation of specific antibiotics detrimental to Xee85-10^25,34,36,40,57–60^.

*Pst*DC3000 growth showed only minimal inhibition by *Bv*FZB42 supernatants under constantly wet conditions, at supernatant concentration of 30% v/v, yet faced significant inhibition under wet-dry cycles. This variation suggests that the antibiotic(s) produced by *Bv*FZB42 affecting *Pst*DC3000 may differ from those inhibiting *Xee85-10*. While literature on *Bv*FZB42 efficacy against *Pst*DC3000 is limited, there is some indication of its potential through specific antimicrobial compounds^61,62^. The mechanism through which *Bv*FZB42’s antibiotics impact *Pst*DC3000 could involve factors other than direct harm to actively growing cells, potentially linked to the drying kinetics observed in wet-dry cycles. During drying solute concentrations, including antibiotics, can increase dramatically due to water evaporation, potentially concentrating the active compounds beyond their minimum inhibitory concentration (MIC) and thereby amplifying their lethal effect^25,40^. Contrary to some antibiotics whose efficacy diminishes under wet-dry conditions, the compound(s) from *Bv*FZB42 targeting *Pst*DC3000 might retain stability and activity, like certain antibiotics previously shown to maintain or even increase effectiveness in such variable hydration environments^40^.

The mutual antagonism between *Bv*FZB42 and *Pst*DC3000, as observed in our study, to the best of our knowledge, has not been reported before. Notably, our experiments demonstrated that *Pst*DC3000 is capable of inhibiting *Bv*FZB42 under both constant wetness and wet-dry cycles (Fig. 4, Supp. Fig. S13). The mechanism underlying the inhibition of *Bv*FZB42 by *Pst*DC3000 is unknown, though there is one study hinting at a potential underlying mechanism involving competitive behaviors linked to siderophore production^63^.

An interesting phenomenon became apparent when both the supernatants and cells of *Bv*FZB42 were present with *Xee*85-10; the detrimental impact on *Xee*85-10 was less pronounced compared to when only *Bv*FZB42 supernatants were used. This pattern persisted under conditions of constant wetness (Fig. 4) as well as during wet-dry cycles (Fig. 6A). Under wet-dry cycle conditions, this phenomenon could be attributed to the enhanced survival of bacteria in larger microdroplets under MSW conditions. Higher local density of bacterial cells, and aggregates in particular, has been shown to lead to the formation of larger microdroplets on drying surfaces^25^. This raises the possibility that the presence of more bacterial cells within the same macro-droplet, regardless of their potential interactions, can improve survival rates on drying surfaces. Although there was a noted reduction in *Bv*FZB42 CFUs under these conditions (wet-dry cycle 1:1 + supernatants) (Fig. 5B), the presence of non-viable cells cannot be discounted. Single-cell microscopy might help to explain the increased survival and assess the proposed mechanism.

Our study provides important insights into microbial ecology in water-unsaturated environments, but it has certain limitations. Our experiments, designed to simulate bacterial life on surfaces undergoing changes in wetness, were conducted using a medium that, while capturing some aspects of these conditions, lacks the complexity found on surfaces in natural microbial habitats^64,65^. Furthermore, while our findings suggest that mechanisms at the microscale, and small census populations within MSW droplets^66^, may offer a potential explanation for some of the observed phenomena, we did not conduct investigations at this level. Identifying and examining these microscale interactions represents an important direction for future research, which could provide valuable insights into the underlying processes. In addition, we note that identifying the exact molecular mechanism and the specific antibiotics that likely drive the inhibition of each pathogen by the *Bacillus*, was not within the scope of this work. Future studies could explore this area.

This study highlights the profound influence of hydration conditions on interspecies bacterial interactions, in particular antibiotic-mediated interference competition. We observed that competition outcome under wet-dry cycles can significantly differ from those in constantly wet environments. These variations are likely influenced by the unique properties of microscopic surface wetness – prevalent on terrestrial microbial habitats – and by its effect on bacterial physiology. Furthermore, the outcome of interactions in droplets may also diverge from those observed in standard lab experiments, whether in liquid suspension or on agar plates. Our research has potential implications for biocontrol applications of plant pathogens, suggesting that the dynamics observed could influence the effectiveness of biological control strategies. It shows that hydration conditions are important, and that optimal hydration conditions can vary depending on the exact agent-pathogen pair. In conclusion, hydration conditions and their dynamics, in particular wet-dry cycles, play a pivotal role in shaping the outcomes of antibiotic-mediated competition among bacteria.

## Materials and Methods

### Bacterial strains and growth conditions

The following bacterial strains were used in this study: *Xanthomonas euvesicatoria pv. euvesicatoria* 85-10, *Pseudomonas syringae* DC3000 (both strains obtained from the late Guido Sessa, *Pst*DC3000), both were transformed with the mCherry fluorescent plasmid pmp7605^67^ (transformation details below). *Bacillus velezenzis* FZB42_FB01 expressing gfp (obtained from Rainer Borriss, AmyloWiki) ^68^.

In the beginning of each experiment, a small portion of the thawed glycerol stock was transferred onto the surface on fresh LB agar + antibiotics plate (1 µg/ml Erytromycin for *B*.*velezensis*, 50 µg/ml Gentamicin sulfate for *Xee*85-10, and *Pst*DC3000). Then a single colony of each strain was inoculated into 5 ml of a fresh LB + appropriate antibiotic medium. Bacterial cultures were grown for 24 hours under agitation set at 220 rpm, at 28 °C. Then, each culture was centrifuged (Eppendorf, centrifuge model. 5430 R, 2935 rcf, 5 minutes), washed, and resuspended in MTG medium (M9×1, Glucose 4%, Tryptone 1%) to reach an OD of 0.1 in 5ml volume. Next, the cultures were grown for another 24 hours under agitation set at 220 rpm, at 28 °C.

Preparation of supernatants of *Bv*FB42, were done in a similar method to the mentioned above. However, after centrifugation, the pellet was resuspended in 20 ml of MTG medium to reach OD of 0.1 and grown for 24 hours in similar conditions. Then, the culture was centrifuged and filtered using a 0.22 um filter in order to separate between the bacteria cells and their secreted products. Centrifugation and filtration were done twice. Supernatants were kept in 4 °C for up to four days.

Each experiment was initiated with bacterial inoculum of ≈ 5 x 10^7^ to 10^8^ CFU/ml.

### Electrophoresis

To transform both *Xee*85-10 and *Pst*DC3000 with pmp7605 plasmid (mCherry+), electrocompetent cells were prepared as described previously^69^. Transformation was performed by electrophoresis (micropulser, BIO-RADUSA, program 1).

### Minimal inhibitory concentration and inhibition zone assays

To evaluate the antibacterial effect of *Bv*FZB42 cells and supernatants on *Xee*85-10 and *Pst*DC3000, Minimal Inhibitory Concentration (MIC) and inhibition zone assays were performed. To perform MIC assay, cells were diluted into 96 microtiter plate (final OD_600_ = 0.06 for *Xee*85-10 and 0.2 for *Pst*DC3000) set with a twofold serial dilution of supernatants whereas the highest concentration of supernatants was 50% v/v. OD_600_ reads were taken every 30 minutes for 24 hours using a plate reader (Synergy H1 Microplate Reader, BioTek Instruments, USA). MIC values were determined by the minimal concentration in which there was no significant bacterial growth. MIC assay was conducted before each experiment in order to validate that different supernatant batches had the same affect. The MIC chosen for all subsequent experiments, which will be described below, corresponds to 2.4 times the MIC determined for *Xee*85-10 (30% v/v).

For the inhibition zone assay, 1 ml of *Xee*85-10 and *Pst*DC3000 cells with an OD_600_ = 0.5 or 1, respectively, that were diluted to 10^−4^, and evenly spread onto 130mm LB-agar plates. After a 1-hour incubation, discs containing either distilled water (DDW), *Bv*FZB42 supernatants or live bacterial cells (OD_600_=0.5) were carefully placed on the inoculated plates. Subsequently, the plates were incubated at 28°C for 3 days, and the effectiveness of inhibition was qualitatively assessed by the presence of a clear area around the discs.

### Monoculture experiment with *Bv*FZB42 supernatants: Wet-dry cycles and constantly wet conditions experimental setup

Monoculture experiments were performed to evaluate the effect of wet-dry cycle on *Xee*85-10 and *Pst*DC3000 response to the supernatants of *Bv*FZB42. Two conditions were investigated in these experiments, the first condition was bacteria without supernatants, as a control, and the second was bacteria (*Xee*85-10 or *Pst*DC3000) with 30%v/v supernatants (30% of the total volume was *Bv*FZB42 supernatants, prepared as described above). In these experiments, droplets of the following volumes: 1 µl, 2 µl,5 µl,10 µl,20 µl and 100 µl were deposited on the center of a well of a glass-bottom 24-well plate (24-well glass-bottom plate #1.5—Cellvis, USA). A total of six repeats (i.e., drops) was used for each tested drop volume. The 24-well plates were incubated in a growth chamber (Aralab, FITOCLIMA 600-PLH) for 24 hours at 25 °C and 85% relative humidity. To induce drying the wet-dry cycles plates were kept without their plastic lid. For constantly wet conditions, the empty cavities between the wells in these plates were filled with 300 µl of H_2_O, and the lid was sealed with tape, to maintain close to 100% relative humidity.

### Drying dynamics experiments

To characterize the drying dynamics of droplets of various volumes (1-100 µl), a microscopy method was employed by capturing time-lapse images of droplets’ area dynamics. Individual droplet was deposited onto the center of a well in a glass-bottom 24-well plate (#1.5 high-performance cover glass, Cellvis, USA), resembling Figure 9A. Three droplets (repeats) were tested for each tested drop volume. The plate, without the lid, was then placed within a stage-top environmental control chamber (H301-K-FRAME, Okolab srl, Italy), pre-equilibrated to a temperature of 28°C and a relative humidity of 75%. Images of the droplets were systematically captured every hour over a total duration of 18 hours. The determination of droplet areas was performed using NIS elements Arv er. 5.03 software.

### Co-culture experiments: Wet-dry cycles and constantly wet conditions experimental setup

Co-culture experiments were performed to evaluate the effect of wet-dry cycle on interference competition between *Bv*FZB42 and *Xee*85-10 or *Pst*DC3000. In this set of experiments, five conditions were investigated, two controls (monocultures of each strain of the pair), 1:1 ratio (*Xee*85-10 or *Pst*DC3000 cells in a similar concentration to *Bv*FZB42 cells), 1:1 ratio with 30% v/v supernatants and monoculture with 30% v/v supernatants. These experiments were conducted with three droplet volumes (2 µl, 10 µl and 100 µl). Droplets were deposited on the center of a well of a glass-bottom 24-well. A total of six repeats (i.e., drops) were used for each volume. Incubation conditions for both wet-dry cycle and constantly wet treatments were identical to the monoculture experiments.

All five conditions were investigated, in parallel, also in well-mixed suspended liquid cultures. The experimental setting mirrored that of the MIC experiments, with similar treatments. For co-cultures plate-reader experiments florescence was measured in addition to optical density (OD). For *Xee*85-10 and *Pst*DC3000, relative fluorescence units (RFU) were measured (mCherry, Em579/Ex616). Similarly, for *Bv*FZB423, RFU was measured (GFP, Ex479/Em520).

### Gradual rewetting protocol

For the rewetting phase of the wet-dry cycle experiments, at time t=24 h the empty cavities between the wells were filled with 300 µl of H_2_O, and the lid was placed on the plate and sealed with a tape. The sealed plate was incubated for 30 min at 28 °C (leading to RH >95% inside the plate). By the end of this step, the ‘dried’ droplets (in MSW form) adsorbed moisture from the humidified air by condensation and deliquescence. At this point, 30 µl of medium was added to each well, then the plates were sealed again and incubated for additional 30 minutes. Finally, medium was added to get a dilution of 10^−1^, 30 µl was taken for serial dilutions and drop assay and the rest was plated on plates with LB agar with appropriate antibiotics. CFU were counted 24 hours or 48 hours after plating for *Bv*FZB42 and *Pst*DC3000, respectively, because each requires a different amount of time to form visible colonies on agar plates.

### Statistical analysis

Comparisons of cell viability (base on CFU counts) among all conditions in wet-dry cycles and constantly wet conditions were conducted using one-way ANOVA, followed by Tukey’s test. The determination of correlations and relationships between droplet size and changes in cell viability (The difference between the number of CFUs at t=24 h and the number of CFUs at t =0 h) was performed using linear regression and Spearman correlation. Six technical replicates were performed for each condition in each experiment. All statistical analysis was performed using GraphPad Prism version 10.2.3 for Windows (GraphPad Software, USA).

## Acknowledgements

ChatGPT was used for assistance in enhancing the readability and language clarity of this manuscript.

## Author contributions

Y.B.M., T.O., and N.K. conceived the study. Y.B.M. and T.O. designed experiments. Y.B.M performed the experiments and performed data and statistical analyses. All authors discussed the results and contributed to the final manuscript. N.K. supervised the project. Y.B.M, T.O., and N.K. wrote the manuscript.

## Data and materials availability

Data and code are available at 10.6084/m9.figshare.25998925. All data needed to evaluate the conclusions in the paper are present in the paper and/or the Supplementary Materials.

