## Supporting Information for "Hydration conditions as a critical factor in antibiotic-mediated bacterial competition outcomes"

**SI Includes:**

1. **Supporting Figures (figures S1-S14)**

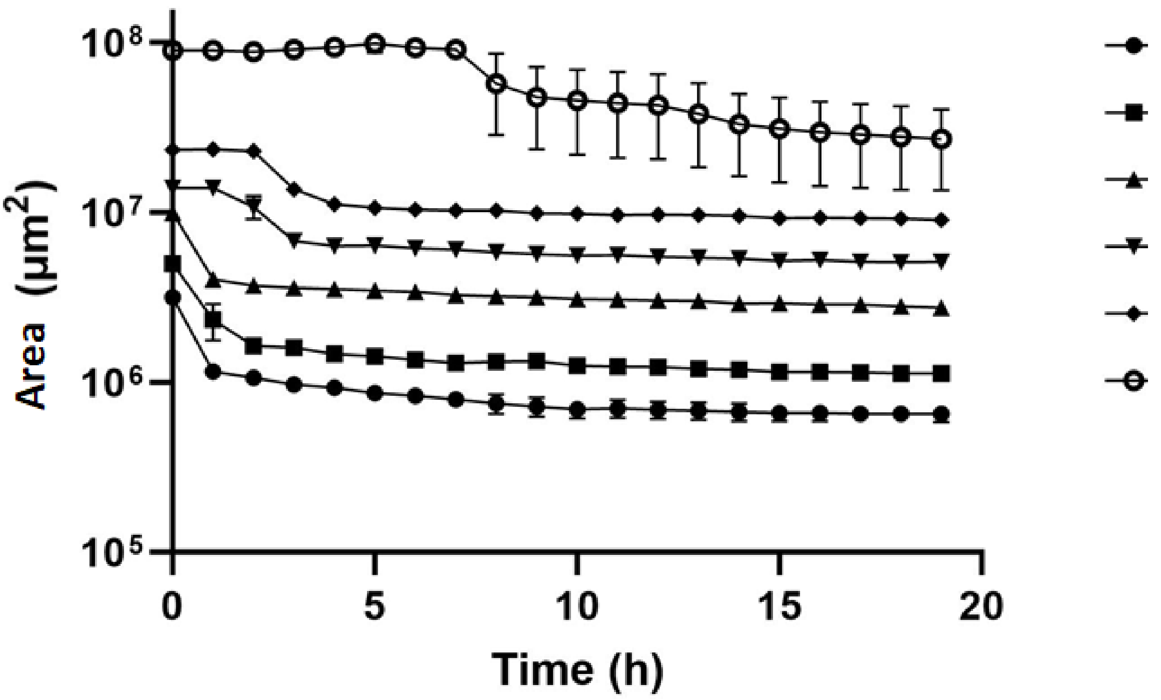


**Fig. S1. Drying dynamics of MTG droplets of various volumes (1 µl to 100 µl).**

Droplets were incubated in 28°C and a relative humidity of 75%. Droplets areas were measured every hour over 18 hours. Each data point and error bar represent the mean ± SEM area (µm²) at specific time points. The experiment details are described in the 'Methods' section.

**Table part 1 — t0 and Constantly wet control**

| | | | Constantly wet control | | | | | |
|---|---|---|---|---|---|---|---|---|
| | | t0 | 1 | 2 | 5 | 10 | 20 | 100 |
| Constantly wet control | 1 | ns | | | | | | |
| | 2 | ns | ns | | | | | |
| | 5 | ns | ns | ns | | | | |
| | 10 | **** | ** | ns | ns | | | |
| | 20 | * | ns | ns | ns | ns | | |
| | 100 | *** | * | ns | ns | ns | ns | |
| Wet-dry cycle control | 1 | * | ** | **** | **** | **** | **** | **** |
| | 2 | ns | ns | *** | **** | **** | **** | **** |
| | 5 | ns | ns | ns | ** | **** | *** | **** |
| | 10 | ns | ns | ns | * | **** | *** | **** |
| | 20 | ns | ns | ns | ns | **** | * | *** |
| | 100 | ns | ns | ns | ns | ns | ns | ns |
| Constantly wet Spn | 1 | **** | **** | **** | **** | **** | **** | **** |
| | 2 | **** | **** | **** | **** | **** | **** | **** |
| | 5 | **** | **** | **** | **** | **** | **** | **** |
| | 10 | **** | **** | **** | **** | **** | **** | **** |
| | 20 | **** | **** | **** | **** | **** | **** | **** |
| | 100 | **** | **** | **** | **** | **** | **** | **** |
| Wet-dry cycle Spn | 1 | **** | **** | **** | **** | **** | **** | **** |
| | 2 | **** | **** | **** | **** | **** | **** | **** |
| | 5 | **** | **** | **** | **** | **** | **** | **** |
| | 10 | **** | **** | **** | **** | **** | **** | **** |
| | 20 | * | ** | **** | **** | **** | **** | **** |
| | 100 | **** | **** | **** | **** | **** | **** | **** |

**Table part 2 — Wet-dry cycle control**

| | | Wet-dry cycle control | | | | | |
|---|---|---|---|---|---|---|---|
| | | 1 | 2 | 5 | 10 | 20 | 100 |
| Constantly wet control | 1 | | | | | | |
| | 2 | | | | | | |
| | 5 | | | | | | |
| | 10 | | | | | | |
| | 20 | | | | | | |
| | 100 | | | | | | |
| Wet-dry cycle control | 1 | | | | | | |
| | 2 | ns | | | | | |
| | 5 | ns | ns | | | | |
| | 10 | ns | ns | ns | | | |
| | 20 | * | ns | ns | ns | | |
| | 100 | **** | *** | ns | ns | ns | |
| Constantly wet Spn | 1 | **** | **** | **** | **** | **** | **** |
| | 2 | **** | **** | **** | **** | **** | **** |
| | 5 | **** | **** | **** | **** | **** | **** |
| | 10 | **** | **** | **** | **** | **** | **** |
| | 20 | ** | **** | **** | **** | **** | **** |
| | 100 | * | **** | **** | **** | **** | **** |
| Wet-dry cycle Spn | 1 | **** | **** | **** | **** | **** | **** |
| | 2 | **** | **** | **** | **** | **** | **** |
| | 5 | *** | **** | **** | **** | **** | **** |
| | 10 | *** | **** | **** | **** | **** | **** |
| | 20 | ns | ns | ns | ns | * | **** |
| | 100 | ns | ** | **** | **** | **** | **** |

**Table part 3 — Constantly wet Spn**

| | | Constantly wet Spn | | | | | |
|---|---|---|---|---|---|---|---|
| | | 1 | 2 | 5 | 10 | 20 | 100 |
| Constantly wet control | 1 | | | | | | |
| | 2 | | | | | | |
| | 5 | | | | | | |
| | 10 | | | | | | |
| | 20 | | | | | | |
| | 100 | | | | | | |
| Wet-dry cycle control | 1 | | | | | | |
| | 2 | | | | | | |
| | 5 | | | | | | |
| | 10 | | | | | | |
| | 20 | | | | | | |
| | 100 | | | | | | |
| Constantly wet Spn | 1 | | | | | | |
| | 2 | ns | | | | | |
| | 5 | ns | ns | | | | |
| | 10 | ns | ns | ns | | | |
| | 20 | *** | *** | ns | ns | | |
| | 100 | **** | **** | ns | * | ns | |
| Wet-dry cycle Spn | 1 | ns | ns | ns | ns | ns | ns |
| | 2 | ns | ns | ns | ns | ns | ns |
| | 5 | ** | ** | ns | ns | ns | ns |
| | 10 | ** | ** | ns | ns | ns | ns |
| | 20 | **** | **** | **** | **** | ** | * |
| | 100 | **** | **** | ** | *** | ns | ns |

**Table part 4 — Wet-dry cycle Spn**

| | | Wet-dry cycle Spn | | | | | |
|---|---|---|---|---|---|---|---|
| | | 1 | 2 | 5 | 10 | 20 | 100 |
| Constantly wet control | 1 | | | | | | |
| | 2 | | | | | | |
| | 5 | | | | | | |
| | 10 | | | | | | |
| | 20 | | | | | | |
| | 100 | | | | | | |
| Wet-dry cycle control | 1 | | | | | | |
| | 2 | | | | | | |
| | 5 | | | | | | |
| | 10 | | | | | | |
| | 20 | | | | | | |
| | 100 | | | | | | |
| Constantly wet Spn | 1 | | | | | | |
| | 2 | | | | | | |
| | 5 | | | | | | |
| | 10 | | | | | | |
| | 20 | | | | | | |
| | 100 | | | | | | |
| Wet-dry cycle Spn | 1 | | | | | | |
| | 2 | ns | | | | | |
| | 5 | ns | ns | | | | |
| | 10 | ns | ns | ns | | | |
| | 20 | **** | **** | *** | *** | | |
| | 100 | ** | ns | ns | ns | ns | |

**Fig. S2. Pairwise comparison of the mean CFU/ml ($Log_{10}$ transformed) for each volume and treatment- *Xanthomonas* monoculture.**

The significance was assessed by one-way ANOVA. Significance marked by *,**, *** or ****, denoting p-values of <0.05, <0.005, <0.0005 or <0.0001, respectively. Data points are same as presented in Fig 2A. Analysis was performed in Graphpad prism.

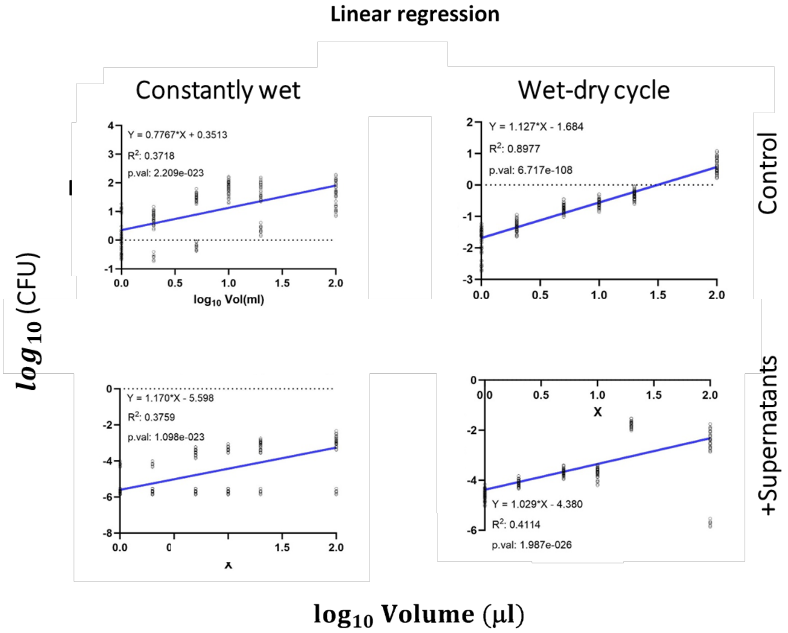


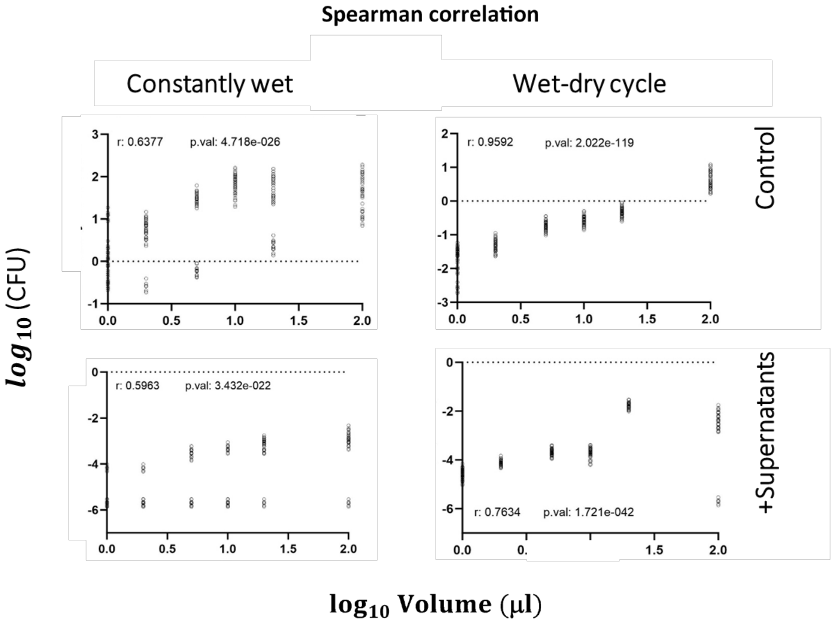


**Fig. S3. Linear regression and spearman correlation of the change in *Xee*85-10 CFU/ml as function of volume, after 24 hours exposure to *Bv*FZB42 supernatants.**

A. linear regression analysis of the change of CFU/ml of *Xee*85-10 cells after exposure of 24 hours to supernatants produced by *Bv*FZB42 in constantly wet and wet-dry cycle conditions. Circles mark experimental values, regression line (blue line) is shown along with its linear equation, $R^2$ values and P-values. B. Correlation between change in bacterial CFU ($Log_{10}$ transformed) and droplet volume ($Log_{10}$ transformed). Circles mark experimental values. r values are Pearson correlation coefficients. Data points are same as presented in Fig. 2B. Analysis was performed in GraphPad Prism

| | | | Constantly wet control | | | | | | Wet-dry cycle control | | | | | | Constantly wet Spn | | | | | | Wet-dry cycle Spn | | | | | |
|---|---|---|---|---|---|---|---|---|---|---|---|---|---|---|---|---|---|---|---|---|---|---|---|---|---|---|
| | | t0 | 1 | 2 | 5 | 10 | 20 | 100 | 1 | 2 | 5 | 10 | 20 | 100 | 1 | 2 | 5 | 10 | 20 | 100 | 1 | 2 | 5 | 10 | 20 | 100 |
| Constantly wet control | 1 | **** | | | | | | | | | | | | | | | | | | | | | | | | |
| | 2 | **** | ns | | | | | | | | | | | | | | | | | | | | | | | |
| | 5 | **** | ns | ns | | | | | | | | | | | | | | | | | | | | | | |
| | 10 | **** | ns | ns | ns | | | | | | | | | | | | | | | | | | | | | |
| | 20 | **** | ns | ns | ns | ns | | | | | | | | | | | | | | | | | | | | |
| | 100 | **** | ns | ns | ns | ns | ns | | | | | | | | | | | | | | | | | | | |
| Wet-dry cycle control | 1 | **** | **** | **** | **** | **** | **** | **** | | | | | | | | | | | | | | | | | | |
| | 2 | **** | **** | **** | **** | **** | **** | **** | **** | | | | | | | | | | | | | | | | | |
| | 5 | **** | **** | **** | **** | **** | **** | **** | **** | ns | | | | | | | | | | | | | | | | |
| | 10 | ns | **** | **** | **** | **** | **** | **** | **** | ** | ** | | | | | | | | | | | | | | | |
| | 20 | ns | **** | **** | **** | **** | **** | **** | **** | **** | *** | ns | | | | | | | | | | | | | | |
| | 100 | ns | * | ** | **** | **** | **** | **** | **** | **** | **** | *** | * | | | | | | | | | | | | | |
| Constantly wet Spn | 1 | ** | ns | ns | * | *** | * | ** | **** | **** | **** | **** | **** | ns | | | | | | | | | | | | |
| | 2 | **** | ns | ns | ns | * | ns | * | **** | **** | **** | **** | **** | ns | ns | | | | | | | | | | | |
| | 5 | *** | ns | ns | ns | ** | ns | * | **** | **** | **** | **** | **** | ns | ns | ns | | | | | | | | | | |
| | 10 | ** | ns | ns | * | *** | * | ** | **** | **** | **** | **** | **** | ns | ns | ns | ns | | | | | | | | | |
| | 20 | ** | ns | ns | * | *** | * | ** | **** | **** | **** | **** | **** | ns | ns | ns | ns | ns | | | | | | | | |
| | 100 | **** | ns | ns | ns | ns | ns | ns | **** | **** | **** | **** | **** | ns | ns | ns | ns | ns | ns | | | | | | | |
| Wet-dry cycle Spn | 1 | **** | **** | **** | **** | **** | **** | **** | ns | **** | **** | **** | **** | **** | **** | **** | **** | **** | **** | **** | | | | | | |
| | 2 | **** | **** | **** | **** | **** | **** | **** | ns | **** | **** | **** | **** | **** | **** | **** | **** | **** | **** | **** | ns | | | | | |
| | 5 | **** | **** | **** | **** | **** | **** | **** | ns | **** | **** | **** | **** | **** | **** | **** | **** | **** | **** | **** | ns | ns | | | | |
| | 10 | **** | **** | **** | **** | **** | **** | **** | ns | **** | **** | **** | **** | **** | **** | **** | **** | **** | **** | **** | ns | ns | ns | | | |
| | 20 | **** | **** | **** | **** | **** | **** | **** | ns | **** | **** | **** | **** | **** | **** | **** | **** | **** | **** | **** | ns | ns | ns | ns | | |
| | 100 | **** | **** | **** | **** | **** | **** | **** | ns | **** | **** | **** | **** | **** | **** | **** | **** | **** | **** | **** | ns | ns | ns | ns | ns | |

**Fig. S4. Pairwise comparison of the mean CFU/ml ($Log_{10}$ transformed) for each volume and treatment - *Pseudomonas* monoculture.**

The significance was assessed by one-way ANOVA. Significance marked *,**, *** or ****, denoting p-values of <0.05, <0.005, <0.0005 or <0.0001 respectively. Data points are same as presented in Fig 3A. Analysis was performed in Graphpad prism**.**

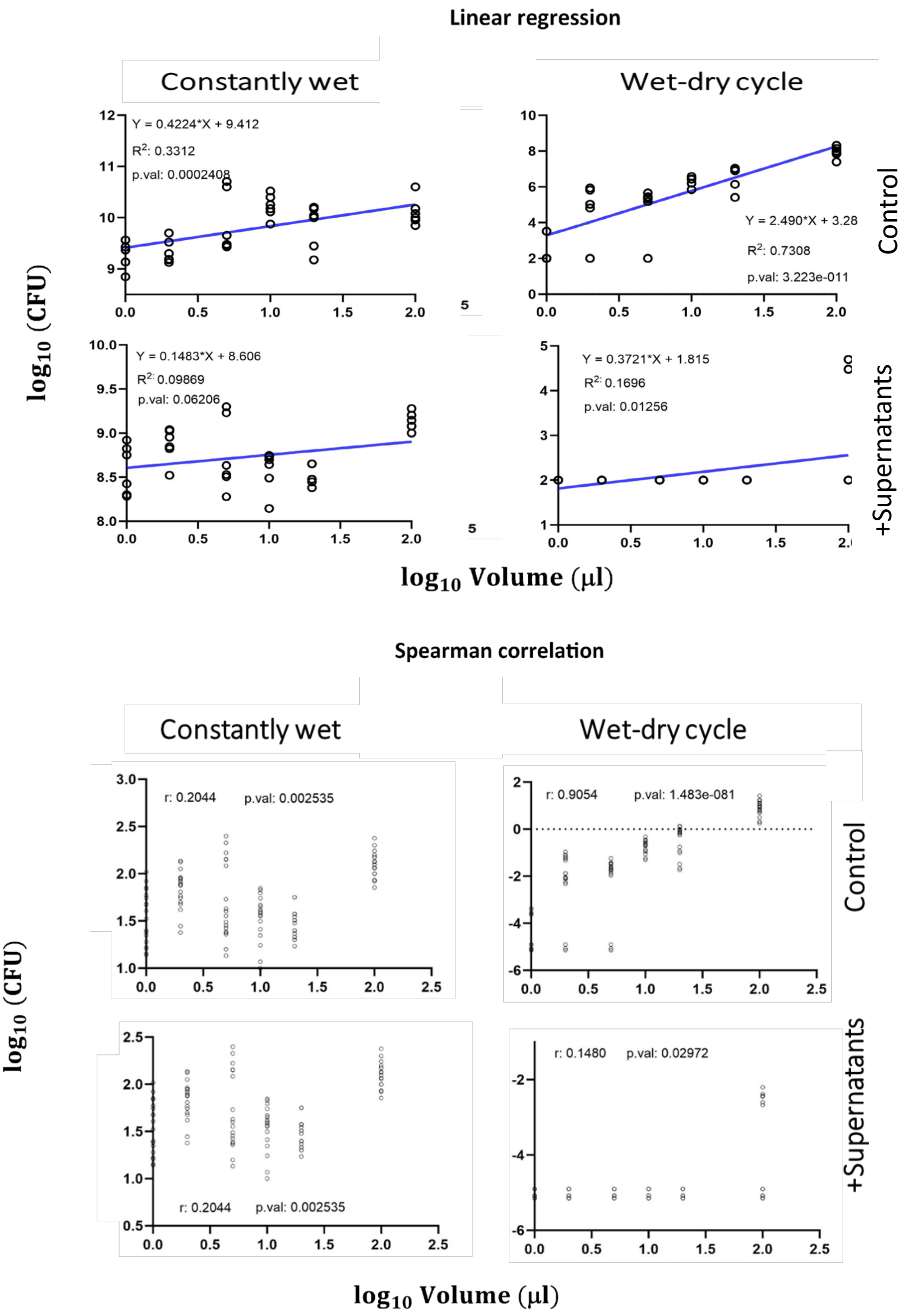


**Fig. S5. Linear regression and spearman correlation of the change in *Pst*DC3000 CFU/ml as function of volume, after 24 hours exposure to *Bv*FZB42 supernatants**.

A. linear regression analysis of the change of CFU/ml of *Pst*DC3000 cells after exposure of 24 hours to supernatants produced by *Bv*FZB42 in constantly wet and wet-dry cycle conditions. Circles mark experimental values, regression line (blue line) is shown along with its linear equation, $R^2$ values and P-values. B. Correlation between change in bacterial CFU ($Log_{10}$ transformed) and droplet volume ($Log_{10}$ transformed). Circles mark experimental values. r values are Pearson correlation coefficients. Data points are same as presented in Fig. 3B. Analysis was performed in GraphPad Prism

| | | | constantly wet control | | | wet-dry cycle control | | | constntly wet 1:1 | | | wet-dry cycle 1:1 | | | constantly wet 1:1+Spn | | | wet-dry cycle 1:1+Spn | | | constantly wet Spn | | | wet-dry cycle Spn | | |
|---|---|---|---|---|---|---|---|---|---|---|---|---|---|---|---|---|---|---|---|---|---|---|---|---|---|---|
| | | t0 | 2 | 10 | 100 | 2 | 10 | 100 | 2 | 10 | 100 | 2 | 10 | 100 | 2 | 10 | 100 | 2 | 10 | 100 | 2 | 10 | 100 | 2 | 10 | 100 |
| constantly wet control | 2 | **** | | | | | | | | | | | | | | | | | | | | | | | | |
| | 10 | **** | ns | | | | | | | | | | | | | | | | | | | | | | | |
| | 100 | **** | ns | ns | | | | | | | | | | | | | | | | | | | | | | |
| wet-dry cycle control | 2 | **** | **** | **** | **** | | | | | | | | | | | | | | | | | | | | | |
| | 10 | **** | **** | **** | **** | ns | | | | | | | | | | | | | | | | | | | | |
| | 100 | ns | **** | **** | **** | **** | **** | | | | | | | | | | | | | | | | | | | |
| constntly wet 1:1 | 2 | ns | **** | **** | **** | * | ns | ns | | | | | | | | | | | | | | | | | | |
| | 10 | ns | **** | **** | **** | **** | **** | ns | ns | | | | | | | | | | | | | | | | | |
| | 100 | ns | **** | **** | **** | ** | ns | ns | ns | ns | | | | | | | | | | | | | | | | |
| wet-dry cycle 1:1 | 2 | ** | **** | **** | **** | ns | ns | ** | ns | * | ns | | | | | | | | | | | | | | | |
| | 10 | **** | **** | **** | **** | ns | ns | **** | *** | **** | *** | ns | | | | | | | | | | | | | | |
| | 100 | **** | **** | **** | **** | ns | ns | **** | **** | **** | **** | *** | ns | | | | | | | | | | | | | |
| constantly wet 1:1+Spn | 2 | **** | **** | **** | **** | * | *** | **** | **** | **** | **** | **** | ns | ns | | | | | | | | | | | | |
| | 10 | **** | **** | **** | **** | **** | **** | **** | **** | **** | **** | **** | *** | ns | ns | | | | | | | | | | | |
| | 100 | **** | **** | **** | **** | ns | * | **** | **** | **** | **** | **** | ns | ns | ns | ns | | | | | | | | | | |
| wet-dry cycle 1:1+Spn | 2 | **** | **** | **** | **** | **** | **** | **** | **** | **** | **** | **** | **** | ** | ns | ns | * | | | | | | | | | |
| | 10 | **** | **** | **** | **** | **** | **** | **** | **** | **** | **** | **** | *** | ns | ns | ns | ns | ns | | | | | | | | |
| | 100 | **** | **** | **** | **** | **** | **** | **** | **** | **** | **** | **** | **** | ** | ns | ns | ns | ns | ns | | | | | | | |
| constantly wet Spn | 2 | **** | **** | **** | **** | **** | **** | **** | **** | **** | **** | **** | **** | **** | **** | *** | **** | ** | **** | ** | | | | | | |
| | 10 | **** | **** | **** | **** | **** | **** | **** | **** | **** | **** | **** | **** | **** | * | ns | *** | ns | ns | ns | ns | | | | | |
| | 100 | **** | **** | **** | **** | *** | **** | **** | **** | **** | **** | **** | ** | ns | ns | ns | ns | ns | ns | ns | **** | ns | | | | |
| wet-dry cycle Spn | 2 | **** | **** | **** | **** | **** | **** | **** | **** | **** | **** | **** | **** | **** | *** | ns | **** | ns | ns | ns | ns | ns | * | | | |
| | 10 | **** | **** | **** | **** | **** | **** | **** | **** | **** | **** | **** | *** | ns | ns | ns | ns | ns | ns | ns | *** | ns | ns | ns | | |
| | 100 | **** | **** | **** | **** | **** | **** | **** | **** | **** | **** | **** | **** | **** | ** | ns | **** | ns | ns | ns | ns | ns | * | ns | ns | |

**Fig. S6. Pairwise comparison of the mean CFU/ml ($Log_{10}$ transformed) for each volume and treatment of *Xee*85-10 in co-culture with *Bv*FZB42.**

The significance was assessed by one-way ANOVA. Significance marked by *,**, *** or ****, denoting p-values of <0.05, <0.005, <0.0005 or <0.0001, respectively. Data points are same as presented in Fig 4A. Analysis was performed in Graphpad prism

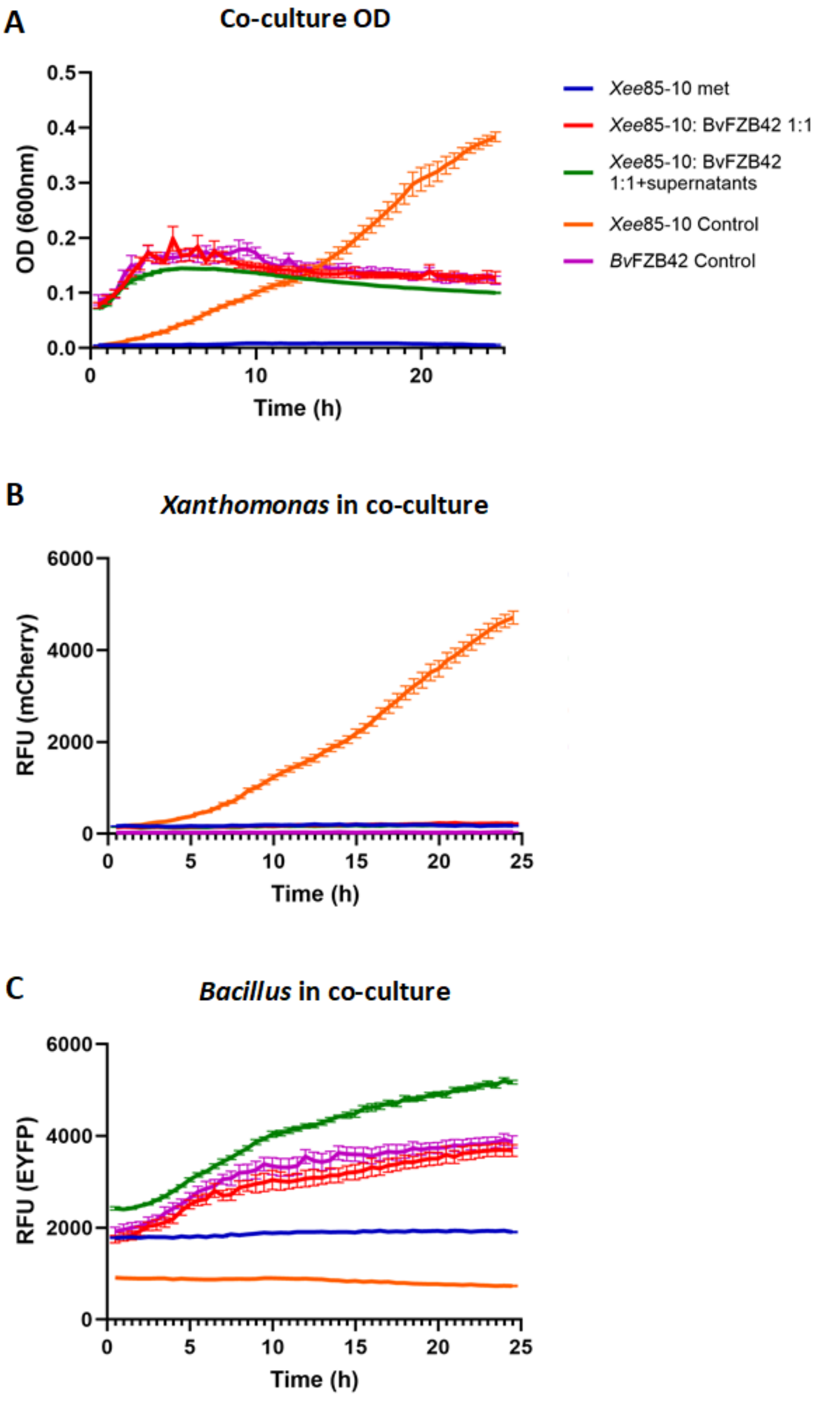


**Fig. S7. Co-culture experiment of *Xee*85-10 and *Bv*FZB42 under continuous shaking.**

A, B and C present the growth curves of the five different treatments in liquid bulk conditions. Four repeats of 200 µl of each treatment were grown in a constantly shaking environment at 28°C in the plate reader (Synergy H1 Microplate Reader, BioTek Instruments, USA). Measurements were taken every 30 minutes for a total of 24 hours. OD (both bacteria) (A), mCherry (Growth of *Xee*85-10 in co-culture) (B), and GFP (growth of *Bv*FZB42 in co-culture) (C) measurements were recorded. Lines and error bars represent mean ± SE.

| | | | constantly wet control | | | wet-dry cycle control | | | constntly wet 1:1 | | | wet-dry cycle 1:1 | | | constantly wet 1:1+Spn | | | wet-dry cycle 1 | |
|---|---|---|---|---|---|---|---|---|---|---|---|---|---|---|---|---|---|---|---|
| | | t0 | 2 | 10 | 100 | 2 | 10 | 100 | 2 | 10 | 100 | 2 | 10 | 100 | 2 | 10 | 100 | 2 | 1 |
| constant ly wet control | 2 | ns | | | | | | | | | | | | | | | | | |
| | 10 | **** | ns | | | | | | | | | | | | | | | | |
| | 100 | **** | ns | ns | | | | | | | | | | | | | | | |
| wet-dry cycle control | 2 | * | **** | **** | **** | | | | | | | | | | | | | | |
| | 10 | ns | ns | **** | **** | ** | | | | | | | | | | | | | |
| | 100 | ns | ns | **** | **** | ns | ns | | | | | | | | | | | | |
| constntl y wet 1:1 | 2 | ns | ns | ns | ns | **** | ns | * | | | | | | | | | | | |
| | 10 | **** | **** | ns | ns | **** | **** | **** | **** | | | | | | | | | | |
| | 100 | **** | ns | ns | ns | **** | **** | **** | ns | ns | | | | | | | | | |
| wet-dry cycle 1:1 | 2 | ns | * | **** | **** | ns | ns | ns | ** | **** | **** | | | | | | | | |
| | 10 | ns | ** | **** | **** | ns | ns | ns | *** | **** | **** | ns | | | | | | | |
| | 100 | ns | ns | ** | ** | **** | ns | ns | ns | **** | ** | ns | ns | | | | | | |
| constant ly wet 1:1+Spn | 2 | *** | ns | ns | ns | **** | ** | **** | ns | * | ns | **** | **** | ns | | | | | |
| | 10 | ns | ns | ns | ns | **** | ns | ** | ns | *** | ns | ** | *** | ns | ns | | | | |
| | 100 | **** | ns | ns | ns | **** | **** | **** | ns | ns | ns | **** | **** | ** | ns | ns | | | |
| wet-dry cycle 1:1+Spn | 2 | **** | **** | **** | **** | **** | **** | **** | **** | **** | **** | **** | **** | **** | **** | **** | **** | | |
| | 10 | ** | **** | **** | **** | ns | *** | * | **** | **** | **** | * | ns | **** | **** | **** | **** | **** | |
| | 100 | **** | **** | **** | **** | **** | **** | **** | **** | **** | **** | **** | **** | **** | **** | **** | **** | * | **** |

**Fig. S8. Pairwise comparison of the mean CFU/ml ($Log_{10}$ transformed) for each volume and treatment of *Bv*FZB42 in co-culture with *Xee*85-10.**

The significance was assessed by one-way ANOVA. Significance marked by *,**, *** or ****, denoting p-values of <0.05, <0.005, <0.0005 or <0.0001, respectively. Data points are same as presented in Fig 4B. Analysis was performed in Graphpad prism.

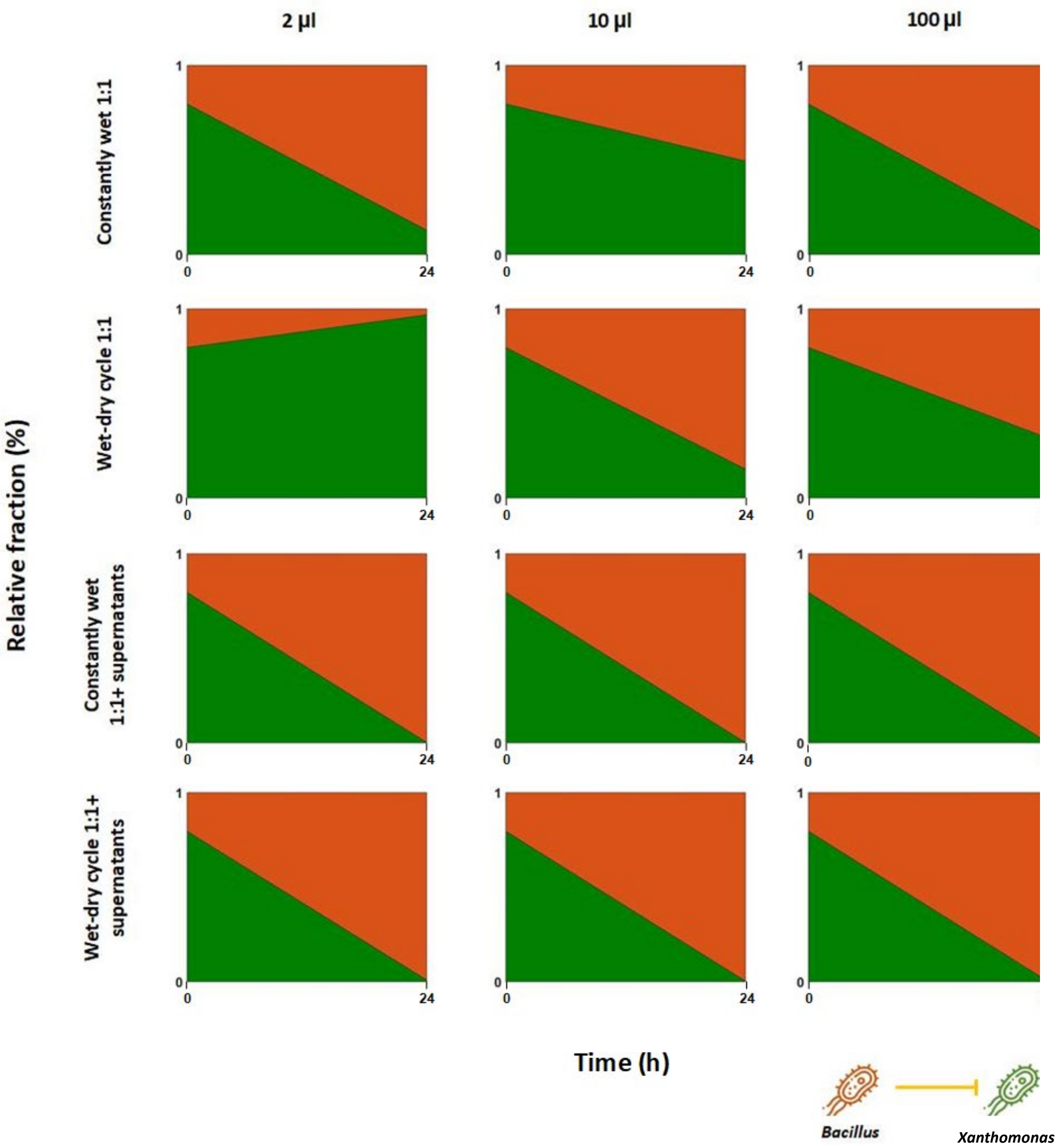


**Fig. S9. Competition dynamics of *Xee*85-10 and *Bv*FZB42 co-culture in droplets.**

The graph shows relative part area plots in droplet co-cultures (1:1 and 1:1+ supernatants). The green fraction represents *Xee*85-10, and the orange fraction represents *Bv*FZB42. Note the plot is based on two time points (at t=0 h and t=24 h).

| | | | constantly wet control | | | wet-dry cycle control | | | constntly wet 1:1 | | | wet-dry cycle 1:1 | | | constantly wet 1:1+Spn | | | wet-dry cycle 1:1+Spn | | | constantly wet Spn | | | wet-dry cycle Spn | | |
|---|---|---|---|---|---|---|---|---|---|---|---|---|---|---|---|---|---|---|---|---|---|---|---|---|---|
| | | t0 | 2 | 10 | 100 | 2 | 10 | 100 | 2 | 10 | 100 | 2 | 10 | 100 | 2 | 10 | 100 | 2 | 10 | 100 | 2 | 10 | 100 | 2 | 10 | 100 |
| constantly wet control | 2 | **** | | | | | | | | | | | | | | | | | | | | | | | | |
| | 10 | **** | ns | | | | | | | | | | | | | | | | | | | | | | | |
| | 100 | **** | ns | **** | | | | | | | | | | | | | | | | | | | | | | |
| wet-dry cycle control | 2 | **** | **** | **** | **** | | | | | | | | | | | | | | | | | | | | | |
| | 10 | ns | **** | **** | **** | ** | | | | | | | | | | | | | | | | | | | | |
| | 100 | **** | *** | **** | ns | **** | **** | | | | | | | | | | | | | | | | | | | |
| constntly wet 1:1 | 2 | * | **** | **** | **** | ns | ns | **** | | | | | | | | | | | | | | | | | | |
| | 10 | * | **** | **** | ** | **** | **** | ns | **** | | | | | | | | | | | | | | | | | |
| | 100 | *** | **** | **** | ns | **** | **** | ns | **** | ns | | | | | | | | | | | | | | | | |
| wet-dry cycle 1:1 | 2 | **** | **** | **** | **** | **** | **** | **** | **** | **** | **** | | | | | | | | | | | | | | | |
| | 10 | **** | **** | **** | **** | **** | **** | **** | **** | **** | **** | **** | | | | | | | | | | | | | | |
| | 100 | **** | **** | **** | **** | **** | **** | **** | **** | **** | **** | **** | ns | | | | | | | | | | | | | |
| constantly wet 1:1+Spn | 2 | **** | **** | **** | **** | **** | **** | **** | **** | **** | **** | ns | **** | **** | | | | | | | | | | | | |
| | 10 | **** | **** | **** | **** | ns | **** | **** | ** | **** | **** | **** | **** | **** | **** | | | | | | | | | | | |
| | 100 | **** | **** | **** | **** | ns | *** | **** | ns | **** | **** | **** | **** | **** | **** | ns | | | | | | | | | | |
| wet-dry cycle 1:1+Spn | 2 | **** | **** | **** | **** | **** | **** | **** | **** | **** | **** | **** | ns | ns | **** | **** | **** | | | | | | | | | |
| | 10 | **** | **** | **** | **** | **** | **** | **** | **** | **** | **** | **** | ns | ns | **** | **** | **** | ns | | | | | | | | |
| | 100 | **** | **** | **** | **** | **** | **** | **** | **** | **** | **** | **** | ns | ns | **** | **** | **** | ns | ns | | | | | | | |
| constantly wet Spn | 2 | **** | *** | **** | ns | **** | **** | ns | **** | ns | ns | **** | **** | **** | **** | **** | **** | **** | **** | **** | | | | | | |
| | 10 | **** | ns | **** | ns | **** | **** | ns | **** | **** | ** | **** | **** | **** | **** | **** | **** | **** | **** | **** | ns | | | | | |
| | 100 | **** | ns | **** | ns | **** | **** | ns | **** | * | ns | **** | **** | **** | **** | **** | **** | **** | **** | **** | ns | ns | | | | |
| wet-dry cycle Spn | 2 | **** | **** | **** | **** | **** | **** | **** | **** | **** | **** | **** | ns | ns | **** | **** | **** | ns | ns | ns | **** | **** | **** | | | |
| | 10 | **** | **** | **** | **** | **** | **** | **** | **** | **** | **** | **** | ns | ns | **** | **** | **** | ns | ns | ns | **** | **** | **** | ns | | |
| | 100 | **** | **** | **** | **** | **** | **** | **** | **** | **** | **** | **** | ns | ns | **** | **** | **** | ns | ns | ns | **** | **** | **** | ns | ns | |

**Fig. S10. Pairwise comparison of the mean CFU/ml ($Log_{10}$ transformed) for each volume and treatment of *Pst*DC3000 in co-culture with *Bv*FZB42.**

The significance was assessed by one-way ANOVA. Significance marked by *,**, *** or ****, denoting p-values of <0.05, <0.005, <0.0005 or <0.0001, respectively. Data points are same as presented in Fig 6A. Analysis was performed in Graphpad prism.

| | | t0 | constantly wet control | | | wet-dry cycle control | | | constntly wet 1:1 | | | wet-dry cycle 1:1 | | | constantly wet 1:1+Spn | | | wet-dry cycle 1:1+Spn | | |
|---|---|---|---|---|---|---|---|---|---|---|---|---|---|---|---|---|---|---|---|---|
| | | | 2 | 10 | 100 | 2 | 10 | 100 | 2 | 10 | 100 | 2 | 10 | 100 | 2 | 10 | 100 | 2 | 10 | 100 |
| constantly wet control | 2 | ns | | | | | | | | | | | | | | | | | | |
| | 10 | ns | ns | | | | | | | | | | | | | | | | | |
| | 100 | ns | ns | ns | | | | | | | | | | | | | | | | |
| wet-dry cycle control | 2 | **** | **** | **** | **** | | | | | | | | | | | | | | | |
| | 10 | ** | **** | **** | **** | ns | | | | | | | | | | | | | | |
| | 100 | **** | **** | **** | **** | ns | ns | | | | | | | | | | | | | |
| constntly wet 1:1 | 2 | **** | **** | **** | **** | ns | ns | ns | | | | | | | | | | | | |
| | 10 | ** | **** | **** | **** | ns | ns | ns | ns | | | | | | | | | | | |
| | 100 | ** | **** | **** | **** | ns | ns | ns | ns | ns | | | | | | | | | | |
| wet-dry cycle 1:1 | 2 | **** | **** | **** | **** | ns | *** | * | ns | ** | ** | | | | | | | | | |
| | 10 | ** | **** | **** | **** | ns | ns | ns | ns | ns | ns | ** | | | | | | | | |
| | 100 | **** | **** | **** | **** | ns | **** | *** | ns | **** | **** | ns | **** | | | | | | | |
| constantly wet 1:1+Spn | 2 | **** | **** | **** | **** | ns | ns | ns | ns | ns | ns | ns | ns | * | | | | | | |
| | 10 | ns | *** | *** | **** | ** | ns | ns | * | ns | ns | **** | ns | **** | ns | | | | | |
| | 100 | ns | ns | ns | ns | **** | **** | **** | **** | **** | **** | **** | **** | **** | **** | **** | | | | |
| wet-dry cycle 1:1+Spn | 2 | **** | **** | **** | **** | ns | **** | ** | ns | **** | **** | ns | **** | ns | * | **** | **** | | | |
| | 10 | **** | **** | **** | **** | ns | ns | ns | ns | ns | ns | ns | ns | ** | ns | ns | **** | ** | | |
| | 100 | **** | **** | **** | **** | ns | ns | ns | ns | ns | ns | ns | ns | ns | ns | ** | **** | ns | ns | |

**Fig. S11. Pairwise comparison of the mean CFU/ml (Log$_{10}$ transformed) for each volume and treatment of *Bv*FZB42 in co-culture with *Pst*DC3000.**

Pairwise comparison of the mean CFU/ml (Log$_{10}$ transformed) of each volume in each treatment. The significance was assessed by one-way ANOVA. Significance marked by *,**, *** or ****, denoting p-values of $<0.05$, $<0.005$, $<0.0005$ or $<0.0001$, respectively. Data points are same as presented in Fig 6B. Analysis was performed in Graphpad prism.

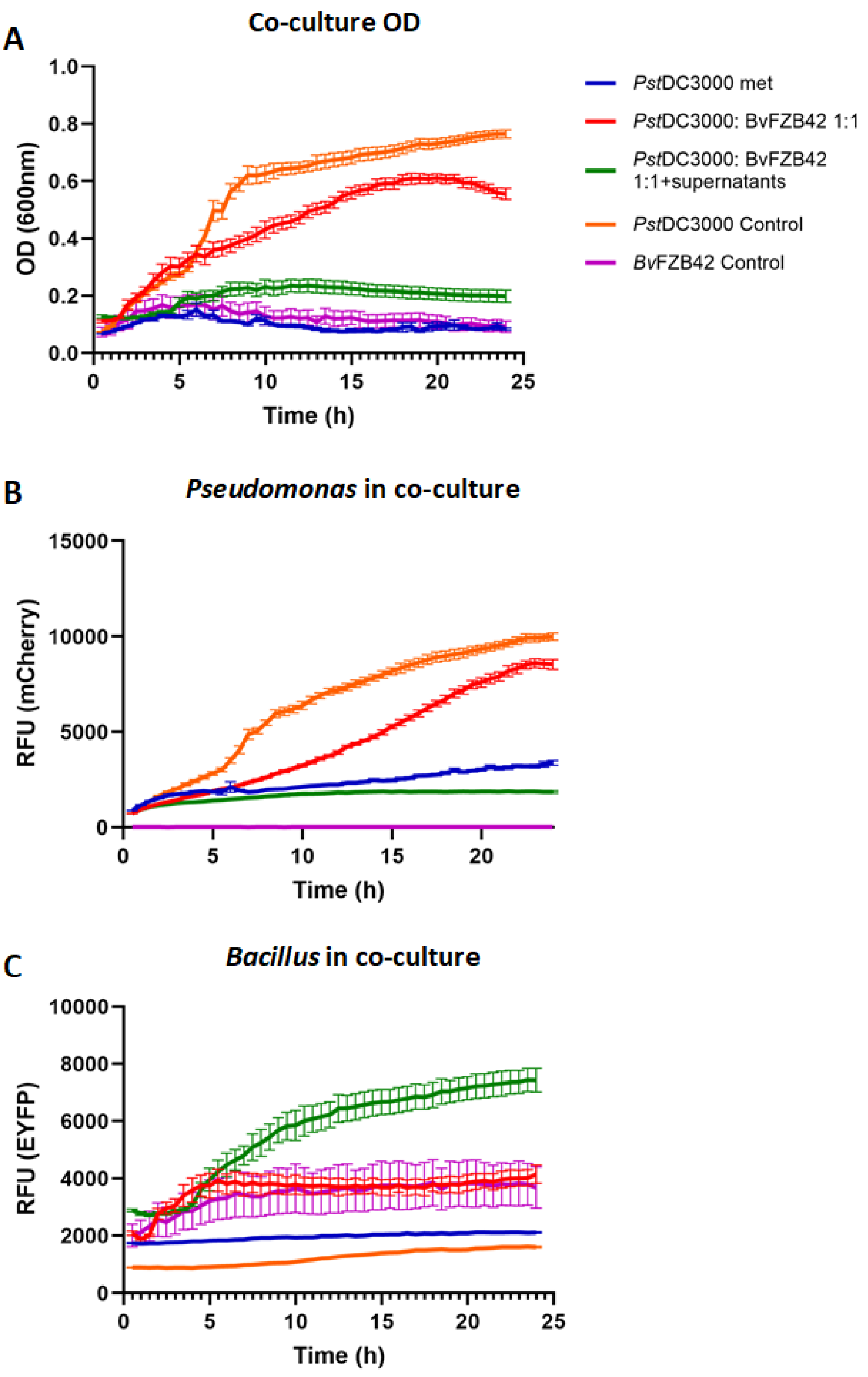


**Fig. S12. Co-culture experiment of *Pst*DC3000 and *Bv*FZB42 under continuous shaking.**

A, B and C present the growth curves of the five different treatments in liquid bulk conditions. Four repeats of 200 µl of each treatment were grown in a constantly shaking environment at 28°C in the plate reader (Synergy H1 Microplate Reader, BioTek Instruments, USA). Measurements were taken every 30 minutes for a total of 24 hours. OD (both bacteria) (A), mCherry (Growth of *Pst*DC3000 in co-culture) (B), and GFP (growth of *Bv*FZB42 in co-culture) (C) measurements were recorded. Lines and error bars represent mean ± SE.

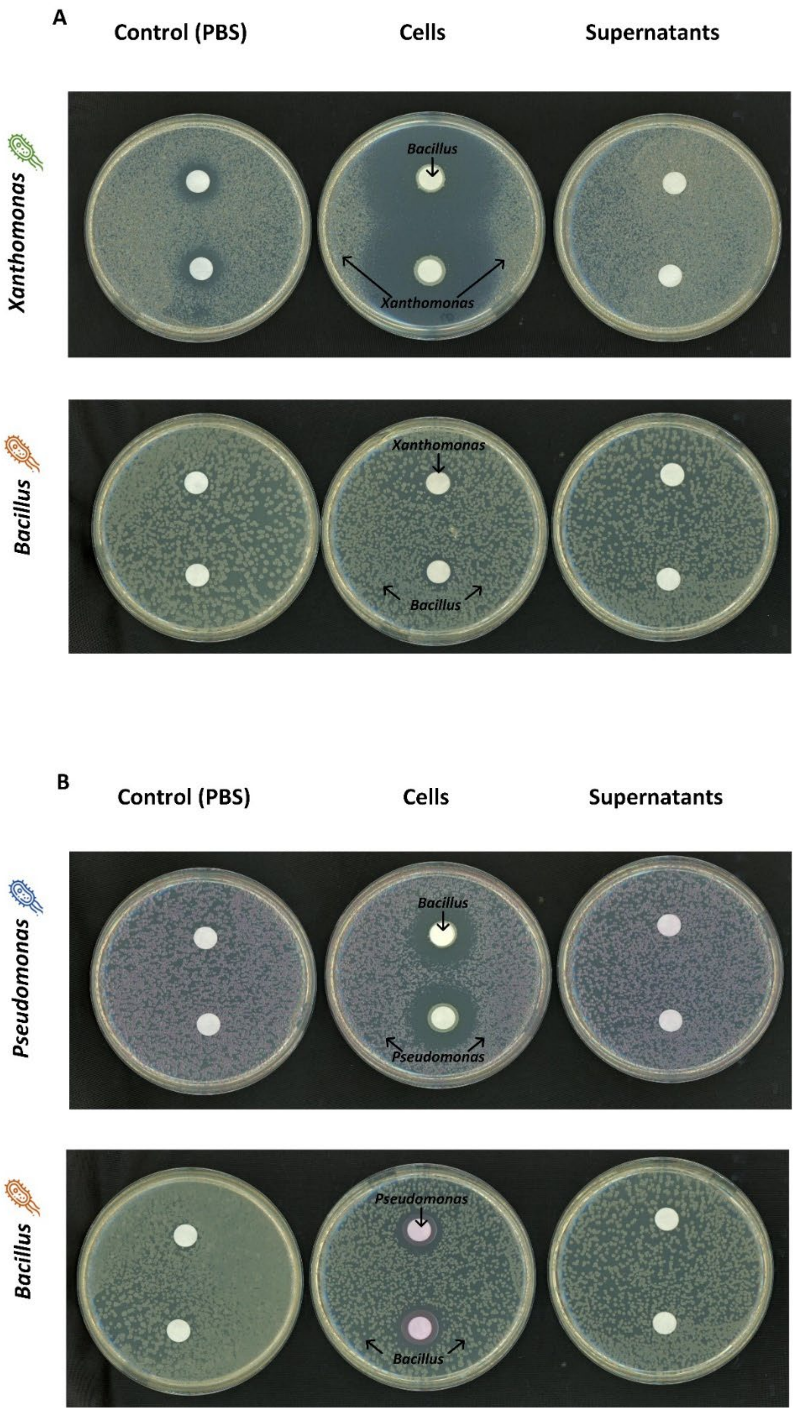


**Fig. S13. Inhibition zone assay of *Xee*85-10 by *Bv*FZB42 and vice versa (A top, bottom, respectively), and *Pst*DC3000 by *Bv*FZB42 and vice-versa (B top, bottom, respectively).**

1 ml of *Xee*85-10, *Pst*DC3000m or *Bv*FZB42 cells with an OD600 = 0.5, 1 or 1, respectively, that were diluted to $10^{-4}$, were evenly spread onto 130 mm LB-agar plates. After a 1-hour incubation, discs containing either distilled water (DDW), supernatants or live bacterial cells (OD600= 0.5) were carefully placed on the inoculated plates. Subsequently, the plates were incubated at 28°C for 3 days, and the effectiveness of inhibition was qualitatively assessed by the presence of a clear area around the discs.

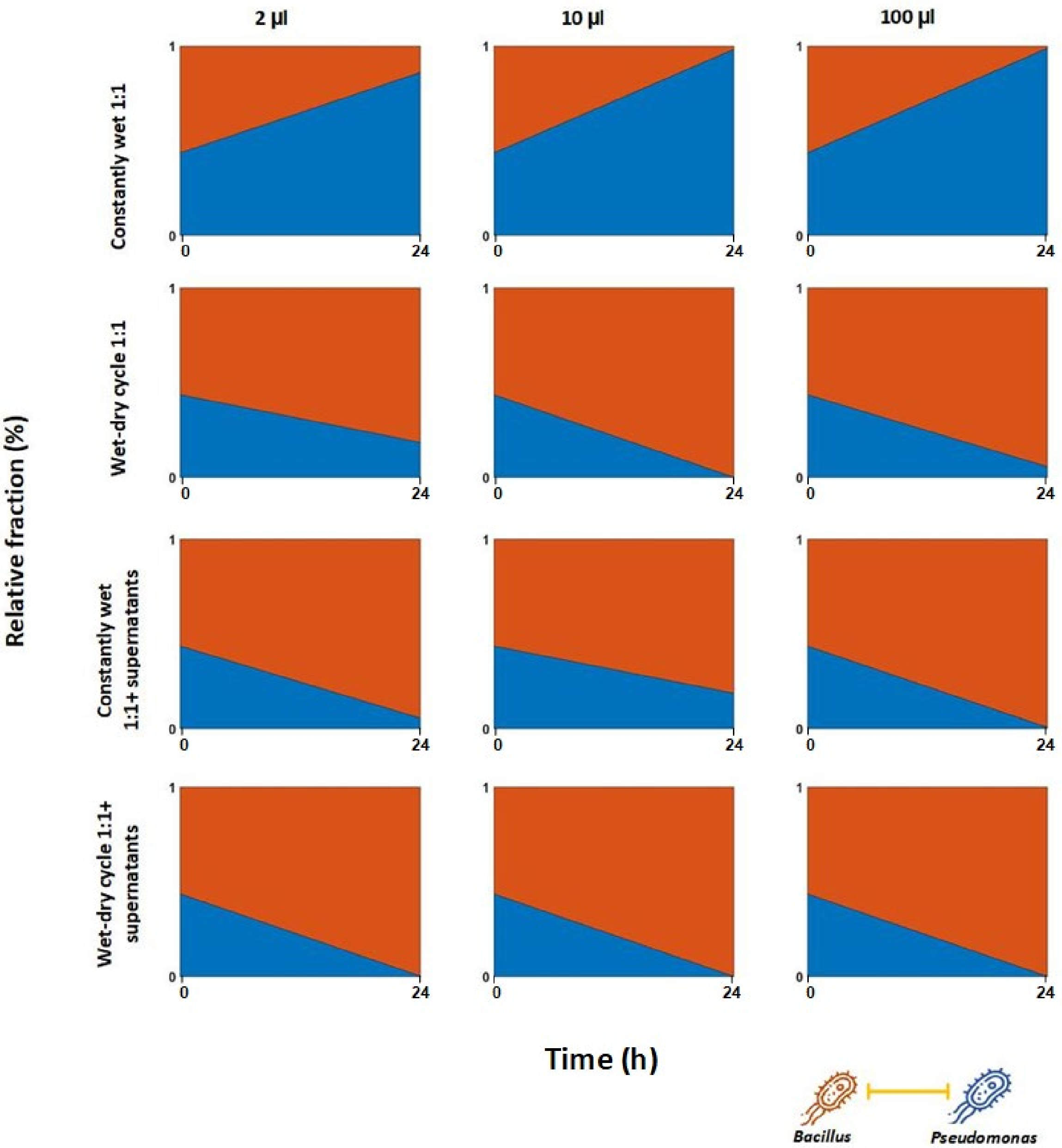


**Fig. S13. Competition dynamics of *Pst*DC3000 and *Bv*FZB42 co-culture in droplets.**

The graph shows relative part area plots in droplet co-cultures (1:1 and 1:1+ supernatants). The blue fraction represents *Pst*DC3000, and the orange fraction represents *Bv*FZB42. Note the plot is based on two time points (at t=0 h and t=24 h).
